# Lysine 27 of histone H3.3 is a fine modulator of developmental gene expression and stands as an epigenetic checkpoint for lignin biosynthesis in Arabidopsis

**DOI:** 10.1101/2022.06.08.495374

**Authors:** Kateryna Fal, Denisa Tomkova, Marie Le Masson, Adi Faigenboim, Emeline Pano, Nickolay Ishkhneli, Netta-Lee Moyal, Claire Villette, Marie-Edith Chabouté, Alexandre Berr, Leor Eshed Williams, Cristel C. Carles

## Abstract

- Chromatin is a dynamic platform within which gene expression is controlled by epigenetic modifications, notably targeting amino acid residues of histone H3. Among them is Lysine 27 of H3 (H3K27), which trimethylation by the Polycomb Repressive Complex 2 (PRC2) is instrumental in regulating spatio-temporal patterns of key developmental genes. H3K27 is also subjected to acetylation, found at sites of active transcription. Most information on the function of histone residues and their associated modifications in plants was obtained from studies of loss-of-function mutants for the complexes that modify them.
- In order to decrypt the genuine function of H3K27, we expressed a non-modifiable variant of H3 at residue K27 (H3.3^K27A^) in Arabidopsis, and developed a multi-scale approach combining in-depth phenotypical and cytological analyses, with transcriptomics and metabolomics.
- We uncovered that the H3.3^K27A^ variant causes severe developmental defects, part of them reminiscent of PRC2 mutants, part of them new. They include early flowering, increased callus formation, and short stems with thicker xylem cell layer. This latest phenotype correlates with mis-regulation of phenylpropanoid biosynthesis.
- Overall, our results reveal novel roles of H3K27 in plant cell fates and metabolic pathways, and highlight an epigenetic control point for elongation and lignin composition of the stem.

## Introduction

Histone proteins, core components of chromatin in Eukaryotes, play key roles in development of multi-cellular organisms by regulating DNA accessibility to the transcriptional machinery. These roles particularly take anchor on Lysine residues of histones that can be dynamically modified by writers, which catalyse different types of post-translational modifications including acetylation and methylation. These modifications can either impact directly DNA-histone interactions (e.g., acetylation) or can be recognized as marks (e.g., methylation) by specific readers, for further regulation of gene accessibility and transcriptional control.

Two key antagonistic complexes, the Polycomb goup (PcG) and trithorax group (trxG), catalyse the deposition of methyl groups on Histone 3 Lysine amino acids. Lysines 4 and 36 (H3K4/K36) can be trimethylated by trxG, forming an active mark for gene transcription, while H3K27 can be trimethylated by the PcG Repressive Complex 2 (PRC2), providing a repressive mark for gene transcription.

Historically, the PcG functions were discovered from mutants in Drosophila that displayed homeotic conversions due to ectopic de-repression of *Hox* genes, while the trxG functions were initially identified from mutants that rescue such PcG phenotypes (Ingham, 1983)(Klymenko & Muller, 2004). Beyond this genetic antagonism, molecular characterization of trxG and PcG functions brought further evidence for their opposed actions on gene transcription (Grimaud *et al*., 2006)(Schuettengruber *et al*., 2007). Moreover, the marks brought at targets by PRC2 and trxG Histone Methyl Transferases (HMTs) are mutually exclusive, through direct prevention of each other’s enzymatic activity (Schmitges *et al*., 2011)(Finogenova *et al*., 2020). PRC2 and trxG activities have counterparts in plants, the PRC2 complex being the best conserved (Engelhorn *et al*., 2014)(Bieluszewski *et al*., 2021)(Baile *et al*., 2022); however, differently from animals, most PRC2 mutants are viable in plants, likely due to redundancies of factors and multiplicity of complexes (Mozgova & Hennig, 2015)(Shu *et al*., 2020)(Bieluszewski *et al*., 2021)(Godwin & Farrona, 2022).

In Arabidopsis, the two sporophytic homologues of the Drosophila E(z) HMT, CURLY LEAF (CLF) and SWINGER (SWN), are responsible for the trimethylation of H3K27 at thousands of target genes, thereby preventing their ectopic transcription (Shu *et al*., 2019). CLF and SWN are partially redundant, having a subset of common target genes predominantly involved in plant growth and development functions, as well as in stimuli responses (Derkacheva & Hennig, 2014)(Shu *et al*., 2019). PRC2 targets are also over-represented with transcription factors. Yet, while *clf* mutants exhibit dwarf stature, curled leaves and early flowering phenotype (Goodrich *et al*., 1997), *swn* mutants display moderate late-flowering phenotypes with no other visible defects (Shu *et al*., 2019)(Shu *et al*., 2020). Loss of both CLF and SWN, though, leads to total loss of body plan and formation of massive somatic embryo-like structures, due to the incapacity of cells to adopt proper fates (Mozgova *et al*., 2017). Loss of function of EMF2, another Arabidopsis PRC2 sub-unit, homologue of the Drosophila Su(z)12 protein and containing a VEFS (VRN2-EMF2-FIS2-Su(z)12) domain, leads to direct flowering upon germination, and sterility (Chanvivattana *et al*., 2004). Thus, despite our knowledge of PRC2 components in plants, the plurality of sub-units, may they be redundant or alternative, does not allow to get a clear idea of the function of the deposited marks themselves (Fal *et al*., 2021).

Compared to HMTs, our knowledge of enzymes responsible for acetylation at H3K27 is sparser, with studies in Arabidopsis reporting GCN5 and TAF1/HAF2 as two histone acetyl transferases, required for H3K9, H3K27, and/or H4K12 acetylation at promoters of Arabidopsis. While GCN5 and TAF1/HAF2 have cumulative effects mainly on H3K9 acetylation, H3K14 acetylation seems to depend only on GCN5 (Benhamed *et al*., 2006). Mutations in *GCN5* and/or *TAF1* cause a long-hypocotyl phenotype and reduced light-inducible gene expression. GCN5 is also involved in cold response, shoot apex identity, leaf cell differentiation, root growth and flower morphogenesis (Bertrand *et al*., 2005) (Bertrand *et al*., 2003) (Poulios & Vlachonasios, 2018) (Servet *et al*., 2010). However, direct target genes of GCN5 and TAF1/HAF2, responsible for these functions, are poorly documented except light-regulated genes (Servet *et al*., 2010) (Bertrand *et al*., 2003).

In summary, most information on the function of histone residues and their associated marks in plants was obtained from studies of loss-of-function mutants in the complexes that modify them. However, such approaches present limits because these complexes are involved into multifaceted interactions, they display redundancies, and are not always specific to a single histone residue. Moreover, total loss of some enzymatic activities (e.g. *clfswn*) causes dramatic phenotypes, preventing to reach a comprehensive functional information.

Several studies in animals (Drosophila and mammals) have explored the effect of mutations at Lysine residues in Histone 3, providing powerful tools to interrogate their roles *in vivo*. Such analyses were extensively done for H3K4, K36 and K27 (Trovato *et al*., 2020). In particular, the Lysine-to-Methionine substitution at K27 shows a dominant phenotype due to its prevailing inhibitory effect on PRC2 HMT (Chan *et al*., 2013)(Herz *et al*., 2014)(Fang *et al*., 2018), thereby contributing to better understand the structural basis of H3K27me3 spreading (Justin *et al*., 2016). Some other studies have used more neutral substitutions with amino acids such as Alanine, revealing interesting effects on gene regulation and development (Pengelly *et al*., 2013)(Leatham-Jensen *et al*., 2019)(Zhang *et al*., 2019)(Gehre *et al*., 2020) (Trovato *et al*., 2020). In contrast, a limited amount of studies reported similar approaches in plants (Iwakawa *et al*., 2017)(Sanders *et al*., 2017)(Lu *et al*., 2018)(Yan *et al*., 2020)(Lin *et al*., 2018) and none of them characterized the effect of a single amino acid substitution at K27 of the ubiquitous H3 proteins, on plant growth and development. Therefore, in order to reveal its true function, we produced *Arabidopsis thaliana* plants that express a variant of Histone 3 lacking this residue (the H3.3^K27A^ variant). We analysed the obtained lines with an integrated strategy, including quantitative phenotyping, histological and cytological analyses, as well as transcriptomics and metabolomics.

We found that developmental and molecular phenotypes of the H3.3^K27A^ Arabidopsis lines recapitulate some of the characteristics reported for loss-of-function in H3K27 modifiers, notably PRC2 mutants. This is particularly true for flowering time and leaf morphology, and the correlated expression of responsible transcription factors. Nonetheless, our dataset also highlighted features unreported thus far, such as strong defects in cell type distribution at the stem, and in related enzymatic pathway components essential for lignin biosynthesis.

## Material and methods

### Plant material and growth conditions

The *Arabidopsis thaliana* plants (Landsberg erecta ecotype, Ler) were grown in growth chambers at 21°C under long-day (LD; 16h/8h light/dark), and for flowering time analyses, as well under medium-day (MD; 12 h light/12 h dark), short-day (SD; 8 h light/16 h dark) or continuous light (CL; 24 h light).

### Construction and selection of transgenic lines

The H3.3 and H3.3^K27A^-encoding DNA fragments were designed from the *HTR5* (At4g40040) gene and obtained by gene synthesis. They include introns, known to be essential to drive the expression of H3.3-encoding genes throughout the cell cycle (Chaubet-Gigot *et al*., 2001). The fragments were inserted into pENTR-D-Topo, transferred into pK2WG7 and transformed into Arabidopsis plants by floral dip (Clough & Bent, 1998). Procedures for plant transformation, primary transformant selection and transgene expression analyses are provided as supporting information (Methods S1).

### Morphological analyses

#### Organogenetic measurements

The lengths of inflorescence stems, quantity of side branches and number of siliques were quantified on plants with fully elongated main stems after all flowers were opened.

Shoot apical meristem sizes were measured on freshly dissected meristems, sampled after the opening of the first flower (Smyth *et al*., 1990). Samples were imaged using a Keyence Digital Microscope VHX-5000 and analysed with FIJI (Schindelin *et al*., 2012), as previously described (Besnard *et al*., 2014).

#### Histological staining for inflorescence stems

Sections (0.5-0.8cm) from the first internode of inflorescence stems from 30-day-old plants were collected, fixed and imbedded in paraplast as previously described (Carles *et al*., 2010)). The toluidine blue and and phloroglucinol stainings were performed as previously described (Pradhan Mitra & Loqué, 2014).

Images were acquired using a Zeiss Imager M2 microscope equiped with an Axiocam 503. All measurements were performed with FIJI. Images of tissue sections stained with phloroglucinol were used for xylem cell size assessment, with the 3D Objects Counter plugin.

#### Statistical analyses

Plots of all presented data sets were prepared using the Rstudio Team (2020) software (http://www.rstudio.com/). The Tukey’s range test was used to make the pairwise comparisons of means from independent samples.

#### Scanning Electron Microscopy (SEM)

SEM analysis was performed as previously described (Talbot & White, 2013) using a JSM-IT100 scanning electron microscope (JEOL Ltd., Japan). In short, fresh tissue was fixed in 100% methanol for 10 min, rinced in 100% ethanol, dried in a K850 critical point dryer (CPD) and mounted on stubs and coated with a thin layer (2 nm) of gold-palladium in a Q150T ES (Quorum Technologies Ltd, UK) for SEM imaging.

### Flowering-time analyses

Analyses were performed as previously described (Berr *et al*., 2015), using two different indicators, the number of days to flowering as a temporal indicator and the leaf number at bolting as a morphometric/developmental indicator.

### Callus formation and regeneration assays

To generate callus, surface-sterilized seeds were sown on MS medium (2% sucrose, 0.8% agar [pH 5.8]) (Murashige & Skoog, 1962) and cultured under CL at 23°C. After 16 days, leaves number 3 and 4 were trimmed and transferred to callus inducing medium (CIM: Gamborg B5 medium 3.2g/L, with 0.5gr/L MES, 2% dextrose, 0.9% phytagel, 2.2μM 2,4-Dichlorophenoxyacetic acid (2,4-D) and 0.46μM kinetin). Plates were placed in the dark, at 23^°^C for seven days, and leaves were further dissected and transferred to a new CIM plate. Calli were re-cultured once a week. All analyses were done on 30-day-old calli.

To test the capacity of 30-day old leaf-derived calli to regenerate shoots, they were transferred to Shoot Inducing Media (SIM: 4.4μM 6-(γ,γ-Dimethylallylamino)purine (2iP) and 0.5μM 1-Naphthylacetic acid (NAA)), placed under continuous light at 23^°^C and imaged using a Leica M205 stereomicroscope.

### Protein extraction and immunobloting

Fractions of soluble/insoluble proteins (Schalk *et al*., 2017) and nuclear protein extracts for histone modification analyses (Zhang *et al*., 2020) were prepared as described previously (see Methods S1 for details). Proteins were separated by 15% SDS-PAGE, transferred onto Immobilon-P membranes (Millipore) using a Trans-Blot cell (Bio-Rad) and analyzed by immunobloting. Antibodies used in this study were anti-UGPase (AS05 086, Agrisera), anti-H3 (C15200011; Diagenode), anti-trimethyl-H3K27 (C15200181; Diagenode), anti-trimethyl-H3K4 (07-473; Millipore); anti-monomethyl-H3K27 (C15410045; Diagenode) or anti-acetyl-H3K27 (C15200184; Diagenode).

### RT-qPCR

RNA was extracted using the Qiagen plant RNeasy Plant Mini Kit (Cat. No. / ID: 74904), and reverse-transcribed to cDNA using SuperScript IV VILO (ThermoFisher, Cat. No. 11756050). The relative transcript abundance was determined using the CYBR Green Master Mix (POWER SYBR GREEN PCR, Thermo Fisher Scientific, 10658255) on a CFX Connect BioRad Real-Time PCR System. Gene-specific primers used for amplification are listed in Table S1.

### RNA-seq for transcriptomic analyses

RNA was extracted from 30-day-old calli, using Qiagen plant RNeasy Plant Mini Kit (Cat.No./ID:74904) and treated with Turbo DNAse kit (Invitrogen, USA). RNA sequencing was performed by Macrogen Inc (Korea) as follow: RNA quality analysis using the 2200 TapeStation System (Agilent Technologies, Santa Clara, CA, USA), mRNA enrichment, library preparation using Illumina TruSeq RNA Library v2 kit, and 50bp single-end sequencing on Illumina NovaSeq. The pipeline for treatment and analysis of raw-reads is provided as supporting information (Methods S1).

Genes with an adjusted *p*-value≤0.05, were considered differentially expressed. Pie charts for Gene Ontology (GO) were drawn from data outputs obtained with the Panther Classification system (http://pantherdb.org). For genes belonging to the “Metabolite Interconversion Enzyme” GO, the KOBAS 3.0 tool (http://kobas.cbi.pku.edu.cn/kobas3/?t=1) and the KEGG PATHWAY module (Kanehisa, 2004) were used to detect statistical enrichments in pathways mapped to the KEGG database (https://www.genome.jp/kegg/).

### Mass-spectrometry for non-targeted metabolomic analyses

Metabolite extraction and analysis was performed on leaf material from 10-day-old seedlings as previously described (Villette *et al*., 2018) using liquid chromatography coupled to high resolution mass spectrometry on an UltiMate 3000 system (Thermo) coupled to an Impact II (Bruker) quadrupole time-of-flight (Q-TOF) spectrometer. Annotations for analyte lists were derived from Phenol Explorer (http://phenol-explorer.eu/), LipidMaps (https://www.lipidmaps.org/), PlantCyc (https://plantcyc.org/), Knapsack (http://www.knapsackfamily.com/). PCA analysis was performed using MetaboAnalyst 5.0. Other statistical analyses were performed in MetaboScape 4.0 with 8 different samples per genotype using the areas of the peaks as the unit of reference. A Wilcoxon rank-sum test was used to compare the different plant extracts and allow the enrichment analysis with a *p*-value set at 0.05 and a minimum fold change at (+/-) 2. The KEGG PATHWAY module (Kanehisa, 2004) was used to identify the enrichment in metabolic pathways.

## Results

### Arabidopsis plants expressing the H3K27A mutant display short stature, early flowering and improved callus production capacity

To address the importance of the Histone 3 Lysine 27 residue in plant development and gene expression, we expressed an artificial H3.3 variant in *Arabidopsis thaliana*, substituting the Lysine residue with an Alanine (H3.3^K27A^) (Fig. 1a-b). We postulated that this strategy should allow to discover novel, genuine functions for the H3K27 residue and its associated marks.

**Figure 1:**
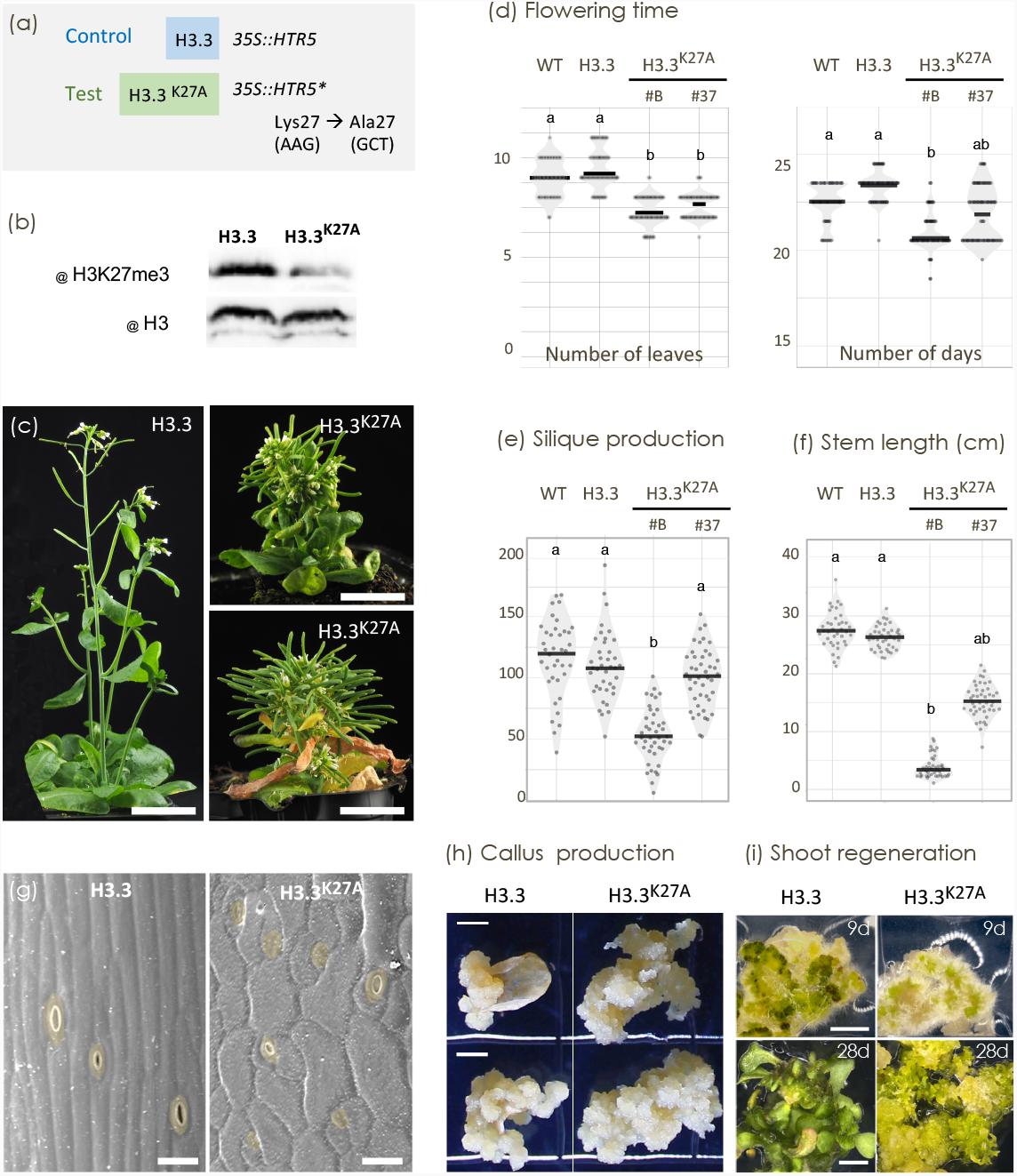
Overview of the alterations in developmental transitions displayed by the *Arabidopsis thaliana* plants expressing H3.3^K27A^: leaf morphology, stem morphology and elongation, flowering transition, flower production, enhanced callus production and impaired shoot regeneration. **(a)** Schematic illustration of the transgenic constructs used to produce the control (H3.3) and test (H3.3^K27A^) lines in the Landsberg erecta (Ler) ecotype. **(b)** Western blot analysis of total H3 accumulation and H3K27me3 level in control H3.3 and H3.3^K27A^ expressing plants. **(c)** Representative images of the adult plant phenotypes from the H3.3 and H3.3^K27A^ lines. Side views of plants at 45 or 60 (right-bottom panel) days post germination. **(d)** Flowering time, scored by number of leaves (left) and number of days (right) at bolting. Plots illustrating the data on the flowering time of the WT, H3.3 and two independent lines of H3.3^K27A^ plants (n = 57, 80, 84 and 87 for WT, H3.3, H3.3^K27A^ #B and H3.3^K27A^ #37, respectively). **(e)** Total number of siliques produced on the main stem (n = 38, 39, 42 and 44 for WT, H3.3, H3.3^K27A^ #B and H3.3^K27A^ #37, respectively). **(f)** Main inflorescence stem length, in cm (n = 38, 39, 42 and 44 for WT, H3.3, H3.3^K27A^ #B and H3.3^K27A^ #37, respectively). Black lines in d to f represent the median and the dots values of individual samples; the samples are assembled in statistical groups by the p-values of Tukey pairwise comparison test. All plants were gown under LD (16h/8h) at 21°C. Data were collected from 3 independently grown plant populations. **(g)** Scanning Electron Microscopy images of inflorescence stem surfaces of 32-day old plants of H3.3 and H3.3^K27A^ genotypes. Stomata cells or founder cells are colorized in yellow. Bar= 25 µm. **(h)** Callus production from leaves 3 and 4 taken from H3.3 and H3.3^K27A^ seedlings and incubated on Callus Inducing Media (CIM) for 30 days. Bar = 200 µm. **(i)** H3.3 and H3.3^K27A^ 30-day old calli were transferred to Shoot Inducing Media (SIM) and images were taken after 9 and 28 days of incubation under light. Bar= 200 µm.

A total of 21 control lines expressing the non-mutated H3.3 (hereafter named H3.3) were undistinguishable from their transformed L*er* wild-type background (WT). On the other hand, 14/31 plants expressing the H3.3^K27A^ variant displayed striking developmental phenotypes, including reduction in stem length and leaf defects, both consistent for all lines (Fig. S2a-b). The stem elongation defects among the 14 H3.3^K27A^ lines could be organised into three classes: no stem elongation (4 lines with stems < 1cm at mature stage; e.g. line #D), short stems (8 lines with stems < 7cm at mature stage; e.g. line #B) and medium-size stems (2 lines with stems <15cm at mature stage; e.g. line #37). Introduction of translationally silent polymorphic sequences in the H3.3-encoding transgenes allowed to identify lines with equivalent level of transgene expression, for comparative analyses between control H3.3 and artificial H3.3^K27A^ variants (Fig. S2c). For all lines, the short stem phenotype intensity correlates positively with the leaf morphology defects -twisted leaves -(Fig. 1c, Fig. S2a-b), as well as with the expression level of the H3.3^K27A^ transgene (Fig. S2c).

As a result of the transgene expression, a substantial amount of unmethylated H3 proteins was detectable in the soluble fraction, for the H3.3 control and H3.3^K27A^ transgenic lines (Fig. S2d). Moreover, as above described for the developmental phenotypes, quantities of soluble H3 proteins correlate positively with the expression level of the transgene (Fig S2c-d). Because phenotypes of the control H3.3 lines were indistinguishable from those of WT plants, the presence of soluble H3 proteins may not be responsible for the severely impaired phenotypes of the H3.3^K27A^ lines. Remarkably, the H3.3^K27A^ transgenic lines, but not the H3.3 control lines, displayed a lower amount of H3K27me3 in the insoluble fraction, expected to correspond to the nucleosome-loaded H3 proteins (Fig. 1b, Fig. S2d).

Expression of the H3.3^K27A^ variant also impacted the floral transition, inducing early flowering, as indicated by numbers of leaves and number of days at bolting (Fig. 1d, Fig. S3). While the control H3.3 line flowered similarly to the WT plants, the H3.3^K27A^ variant provoked an early-flowering phenotype, yet less severe than the PRC2 mutant *clf-2* (Fig. S3) (Carles & Fletcher, 2009). Here again, the severity of the flowering phenotype correlates with the expression level of the mutated H3.3 transgene (Fig. S3). Early flowering could be observed in SD, MD and LD conditions for the H3.3^K27A^ lines, but not under CL (Fig. 1d, Fig. S3). This indicates that both tested H3.3^K27A^ lines retained a photoperiodic response. Supporting the flowering phenotypes, the transcript levels of the flowering repressors *MAF1, MAF2* and *FLC* (Ratcliffe *et al*., 2003) were similarly reduced in both H3.3^K27A^ lines and in *clf-2*, as compared to the control H3.3 line and WT plants (Fig. S4). However, while the transcript levels of both *FT* and *SOC1* floral integrators (Ratcliffe *et al*., 2003) were highly increased in *clf-2, SOC1* appeared more upregulated than *FT* in the two H3.3^K27A^ lines. This latest observation indicates that, comparatively to *clf-2*, early flowering of the H3.3^K27A^ lines is associated more with *SOC1* than with *FT* upregulation. Together, our data point toward multiple similarities between the H3.3^K27A^ lines and the PRC2 HMT mutant *clf-2* in the regulation of flowering genes, yet with some divergences.

Besides being a flowering inducer (Lee *et al*., 2008), *SOC1* is also necessary to limit premature floral meristem differentiation (Lee & Lee, 2010) and cambium activity (Rahimi *et al*., 2022). Interestingly, *SOC1* remains up-regulated in H3.3^K27A^ lines after flowering inducing molecular events have taken place (Fig. S4b). Therefore, to assess organogenesis from the shoot apex, the number of flowers produced by the primary inflorescence and the meristem size, two negatively correlated features (Landrein *et al*., 2015), were measured. While a slight trend towards slower plastochrone could be observed in the H3.3^K27A^ lines (Fig. 1e, Fig. S5), they displayed similar meristem radius and number of side branches as the wild-type control. This indicates that shorter stem length of the H3.3^K27A^ lines is not due to early meristem termination nor loss of apical dominancy (Fig. 1f, Fig. S5).

Finally, the H3.3^K27A^ lines showed higher proliferation rate and mass accumulation upon culturing on callus inducing media (Fig. 1h). Since the PRC2 mutant *emf2* callus displays impaired capacity to regenerate shoot (Mandel *et al*., 2022), we assessed this capacity for 30-day-old H3.3^K27A^ calli, by culturing them on Shoot Inducing Medium (SIM). After 9 days of incubation, the control H3.3 line showed dark green foci characteristic of *de novo* shoot organogenesis initials, whereas many root hairs appeared on H3.3^K27A^ line calli. After 28 days of incubation on SIM, the control H3.3 calli regenerated four to eight shoots whereas the H3.3^K27A^ line calli failed to regenerate ordinary and functional shoots (Fig. 1i-j, Fig. S6). Instead, the calli developed aberrant structures, reminiscent of the PRC2 *clfswn* double mutant (Chanvivattana *et al*., 2004).

### Cell fates are affected in stems of the H3.3^K27A^ lines

The short-stemmed H3.3^K27A^ plants also displayed morphologic defects at the epidermis. Indeed, SEM analysis on stems revealed cells with aberrant shapes, as well as wrong positioning and spacing of stomata initiator cells and incomplete differentiation of stomata from meristemoid mother cells (Fig. 1g, Fig. S7). In order to further characterize morphological defects at inflorescence stems, we performed in depth histological analyses on stem longitudinal and cross sections. We found that the cell type distribution is affected in H3.3^K27A^ stems (Fig. 2a-c). In particular, toluidine blue and phloroglucinol staining of the tissue sections revealed a thickening of the lignified tissue layers with a wide range of increase, from 50% to 100% as compared to the H3.3 control, and an irregularity in distribution of the vascular bundles (Fig.2d, Fig. S8). Further quantification showed that thickening of the scherenchyma layer (xylem and interfascicular fibers) was preferentially due to a significant increase in cell layer number than changes in cell sizes (Fig. 2d; Fig. S8, S9). At the same time, the thickness of the phloem layer appeared unchanged in H3.3^K27A^ stems (Fig. S9). Finally, a tendency for epidermis and cortex layer thickening was also observed in H3.3^K27A^ stems (Fig. S9). These defects, while observed in both analysed transgenic lines, were more dramatic in the globally more affected line #B (Fig. 2; Figs. S8 and S9), once again revealing a positive correlation between strength of phenotypes within a same line. Interestingly, we found that the expression level of the transcription factor *WUSCHEL-related HOMEOBOX4 (WOX4)*, a key regulator of cambium cell division and maintenance of the vascular meristem organization during secondary growth (Hirakawa *et al*., 2010), was reduced in the stems of H3.3^K27A^ plants (Fig. 2e, Fig. S10). A decrease in *WOX4* expression is known to promote xylem differentiation (Hu *et al*., 2022), as observed in the stems of H3.3^K27A^ lines.

**Figure 2:**
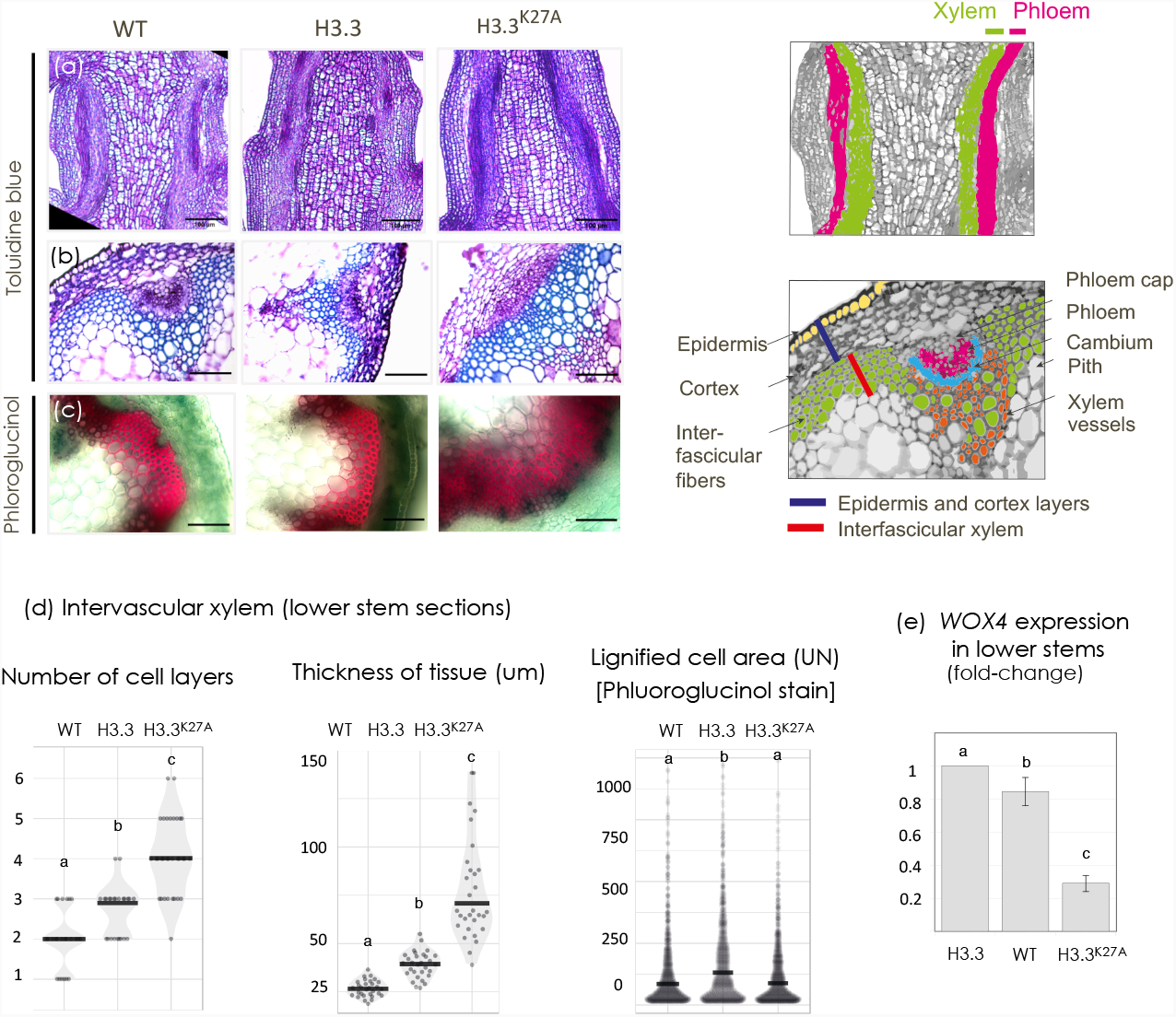
Differentiation of cell layers is affected in stems of the H3.3^K27A^ lines. **(a-b)** Representative images of longitudinal **(a)** and transversal **(b)** microtome sections of paraplast-embedded tissues from the base of the main inflorescence stem of WT, H3.3 and H3.3^K27A^ plants, stained with Toluidine blue. **(c)** Representative images of transversal free hand sections of lower inflorescence stems from H3.3 and H3.3^K27A^ plants (0,7-1 cm above the rosette and under the first side branch, 2.5 weeks after bolting), stained with Phloroglucinol-HCl. Scale bars = 100µm. The right panel represents the schematic drawing of the longitudinal (top) and transversal (bottom) sections of the inflorescence stems with the tissues (color code) and regions taken to measure layer thickness. Scale bars = 100µm. **(d)** Plots illustrating the measurements of the number of cell layers (left panel, n= 24, 24 and 32 for WT, H3.3 and H3.3^K27A^, respectively); thickness of tissue (center, n= 30 for all genotypes); areas of lignified cells (right, n = 1,000 cells for all genotypes). Samples are assembled in statistical groups by the p-values of Student pairwise comparison test. **(e)** Histogram illustrating the abundance of the *WOX4* transcript in the base parts of inflorescence stems of the H3.3, WT and H3.3^K27A^ plants, as measured by RT-qPCR. Data was normalized to *TUB4* and presented relative to H3.3 (set as 1). Error bars represent the standard deviation of the 3 technical repeats, from 1 representative biological repeat; the samples are assembled in statistical groups by the p-values of Student pairwise comparison test. Transgenic lines shown are: H3.3 #C, H3.3^K27A^ #B.

In summary, the K27 of H3.3 plays an important role in leaf morphology, stem elongation, floral transition and *de novo* organogenesis. Moreover, the H3.3^K27A^ lines display phenotypes never reported for the PRC2 mutants, thus revealing novel functions.

### The global level of H3K27me3, but not H3K27me1 is clearly reduced in H3.3 ^K27A^ expressing lines

Next, we analysed the global nuclear and chromatin organization, as well as the histone mark contents of the H3.3^K27A^ lines, by cytological and western blot analyses, respectively.

DAPI Staining and immuno-detection analyses performed *in cyto* did not reveal any clear difference in the nuclear organization and in various nuclear parameters, including distribution of ploidy levels, between H3.3 control and H3.3^K27A^ mutant lines (Figs. S11 and S12).

While the spatial distribution of heterochromatic (H3K9me2, H3K27me1) and euchromatic (H3K4me3, H3K27me3) histone marks did not appear affected in the H3.3^K27A^ mutant lines (Fig. S12), western blots analyses revealed a decrease in H3K27me3 (Figs. S2 and S13). This H3.3^K27A^ variant-induced decrease reached a level similar to that observed in the *clf* mutant (Figs. S2 and S13). Interestingly, the level of H3K27me1, a typical mark of constitutive heterochromatin, appeared unchanged in the H3.3^K27A^ lines, indicating a H3 variant sorting for the mono-methylation of K27. This is in agreement with the report that H3.3 and H3K27me1 are mutually exclusive (Sequeira-Mendes *et al*., 2014). In addition, while *clf-2* loss-of-function causes an increase in the level of H3K27 acetylation, this is not the case for the H3.3^K27A^ lines, in which a slight decrease in H3K27ac is observed, very likely due to the presence of some H3.3 proteins not modifiable on K27. Finally, the opposite effect on the H3K4me3 mark observed in the *clf-2* mutant background was not observed in the H3.3^K27A^ lines, reinforcing the advantage of our approach for studying the genuine function of K27 and associated marks, and avoid collateral effects of the loss of PRC2 activity (Fig. S13).

### RNA-seq on the H3.3^K27A^ plants reveals effects on gene-specific transcriptional regulators and metabolite interconversion enzymes

To study the effect of H3K27A on gene expression, we performed mRNA-seq analysis on 30-day-old calli generated from the H3.3 (#C) control and H3.3^K27A^ (#B) lines. Using calli allowed to compare cells of similar identity, thereby limiting the eventual bias induced by comparing organs of different developmental stages and/or morphologies.

More genes were found to be up-regulated (1764, log2FC≥1) than down-regulated (1337, log2FC≤-1) in calli expressing H3.3^K27A^ as compared to those expressing the control H3.3 (Fig. 3a, Table S2).

**Figure 3:**
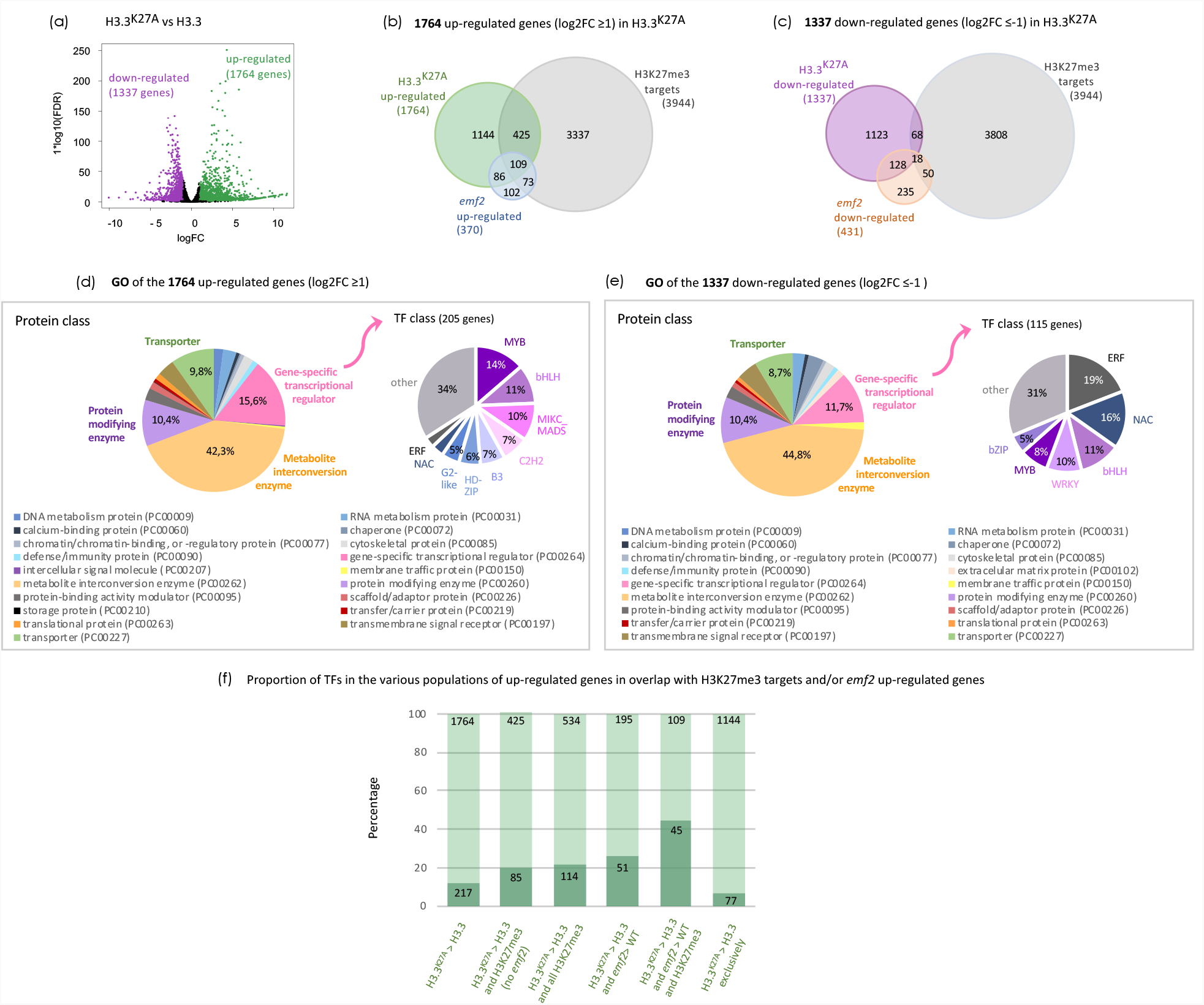
Transcriptomics of the H3.3^K27A^ lines reveal a subset of up-regulated genes larger than that of down-regulated genes, with enrichment in metabolite reconversion enzymes and gene-specific transcriptional regulators. Data analysis of RNA-seq on calli from H3.3^K27A^ vs H3.3 lines. **(a)** Volcano plots for down-and up-regulated genes in H3.3^K27A^ #B compared to H3.3 #C. While 1337 genes are down-regulated (log2FC≤-1 and FDR <0.05), 1764 are upregulated in H3.3^K27A^ (log2FC≥1 and FDR <0.05). **(b**,**c)** Venn diagrams for overlaps between H3.3^K27A^ mis-regulated genes and H3K27me3 targets and/or *emf2* mis-regulated genes. The full list of corresponding genes can be found in the Suppl. Table S2. **(d**,**e)** GO analysis of the genes mis-regulated in H3.3^K27A^, and enrichment in transcription factor (TF) classes (p-value < 0.05). **(f)** percentage of TFs in the various populations of up-regulated genes in overlap with H3K27me3 targets and/or *emf2* up-regulated genes. The full list of corresponding genes can be found in the Suppl. Table S3.

Out of the 1764 up-regulated genes in H3.3^K27A^, 534 (30 %) were identified as H3K27me3-marked genes in WT calli grown under the same conditions (Mandel *et al*., 2022) (Representation factor (RF)=2.1, p-value<8.495e-71 for significant overlap). The up-regulation of these 534 genes is therefore likely a direct consequence of the K27A substitution and the resulting decrease in H3K27me3 (Fig. 3b, Table S3). Furthermore, among the 534 genes up-regulated in H3.3^K27A^, only 195 are part of the 370 *emf2* up-regulated genes (compared with WT, log2FC≥1) (Mandel *et al*., 2022). This indicates that our approach results in a widespread H3K27me3 decrease that goes beyond that of the PRC2 mutant *emf2*, whose smaller population of mis-regulated genes likely reflects redundancy between PRC2 components. Gene Ontology (GO) analyses on the 1764 up-regulated genes in H3.3^K27A^ (log2FC≥1) revealed a large proportion of metabolites interconversion enzymes (42,3%) and gene-specific transcriptional regulators (15,6%) (Fig. 3d). Altogether there are 213 TF-encoding genes up-regulated in H3.3^K27A^, with a majority of helix-turn-helix (HTH), MADS and basic helix-loop-helix (bHLH) encoding genes (Fig. 3d). Interestingly, the proportion of Transcription Factor (TF)-encoding genes increases among the H3.3^K27A^ up-regulated genes covered by H3K27me3 in WT callus (Fig. 3f, Fig. S14). This proportion further increases when considering the 109 genes that are also up-regulated in the *emf2* mutant (Fig. 3d, Fig. S14), especially for MADS genes (from 26.9% to 41.2%).

Remarkably, out of the 431 *emf2* down-regulated genes (log2FC≤-1), 146 genes are also down-regulated in H3.3^K27A^, (Fig. 3c, Table S4) which corresponds to a significant overlap in target lists (RF= 6.9, p-value < 8.891e-83). Because this population of 146 genes, as well as the population of 1337 genes down-regulated in H3.3^K27A^, is independent from the population of H3K27me3 targets (Fig. 3c, Table S4), they likely correspond to secondary targets affected by the up-regulation of PRC2 direct targets, identified in both *emf2* and H3.3^K27A^ lines. The 1337 down-regulated genes in H3.3^K27A^ are again mainly enriched in metabolites interconversion enzymes (44,8%) and gene-specific transcriptional regulators (11,7%). Among the 112 TF-encoding genes that are down-regulated in H3.3^K27A^, 75% belong to the HTH and bHLH families (Fig. 3e), only one gene belongs to the MIKC type MADS box family and 22 to the ERF family (compared with 21 and 6 respectively, in the up-regulated genes) (Fig. 3e, Fig. S15, Table S4).

Interestingly, we found that *SOC1* is highly up-regulated in H3.3^K27A^ calli (log2FC=2,95) (Table S2). This is consistent with our RT-qPCR analyses performed on seedlings (Fig. S4) and further validates our RNA-seq approach on calli. Interestingly, *SOC1* gene is covered by H3K27me3 but not found among the *emf2* up-regulated genes (Mandel *et al*., 2022).

While searching for mis-regulated genes in the H3.3^K27A^ lines, that may be associated with the stem morphology phenotypes, we found *WRKY13* (up-regulated, log2FC= 5,22), previously reported as required for the development of sclerenchyma cells (Table S3). Coherently, the *wrky13* mutant phenotype is opposite to that of the H3.3^K27A^ lines (Li *et al*., 2015). Also related to this phenotype, a considerable proportion of genes mis-regulated in H3.3^K27A^ are associated with metabolic pathways involved in cell wall formation and lignification (Fig. 4d, Table S2), indicating further links with the H3.3^K27A^ stem phenotype. Among them are genes encoding for callose synthase-like protein (At2g30680, log2FC=9.61), cellulose synthase-like B1 (At2g32610, log2FC=6,17), flavonol synthases (up to log2FC=8.36) and peroxidases (expression changes raise up to log2FC=7,43) and ferulic acid 5-hydroxylase 1 (At4g36220, log2FC=1.99) (Table S2). In addition, several TF-encoding genes involved in the regulation of xylem development are associated with H3K27me3-enriched regions, including the bHLH protein TARGET OF MONOPTEROS5 (TMO5) and NAC-domain proteins such as members of the VASCULAR RELATED NAC DOMAIN (VND) group (Hussey *et al*., 2017). These TFs are positive regulators of secondary cell wall formation (Kubo *et al*., 2005)(Ohashi-Ito *et al*., 2010)(Taylor-Teeples *et al*., 2015)(Yamaguchi *et al*., 2011). In particular, overexpression of *VND4* causes ectopic secondary cell wall growth. Coherently with the H3.3^K27A^ stem phenotype, the expression of *VND4* (AT1G12260, log2FC=2.32) and *TMO5* (AT3G25710, log2FC=3.01) is upregulated in H3.3^K27A^ (Tables S2 and S3). Furthermore, our RNA-seq data also reveals the downregulation of *WOX4* in H3.3^K27A^ calli, as previously detected by RT-qPCR on H3.3^K27A^ stem tissues (Tables S2 and S4).

**Figure 4:**
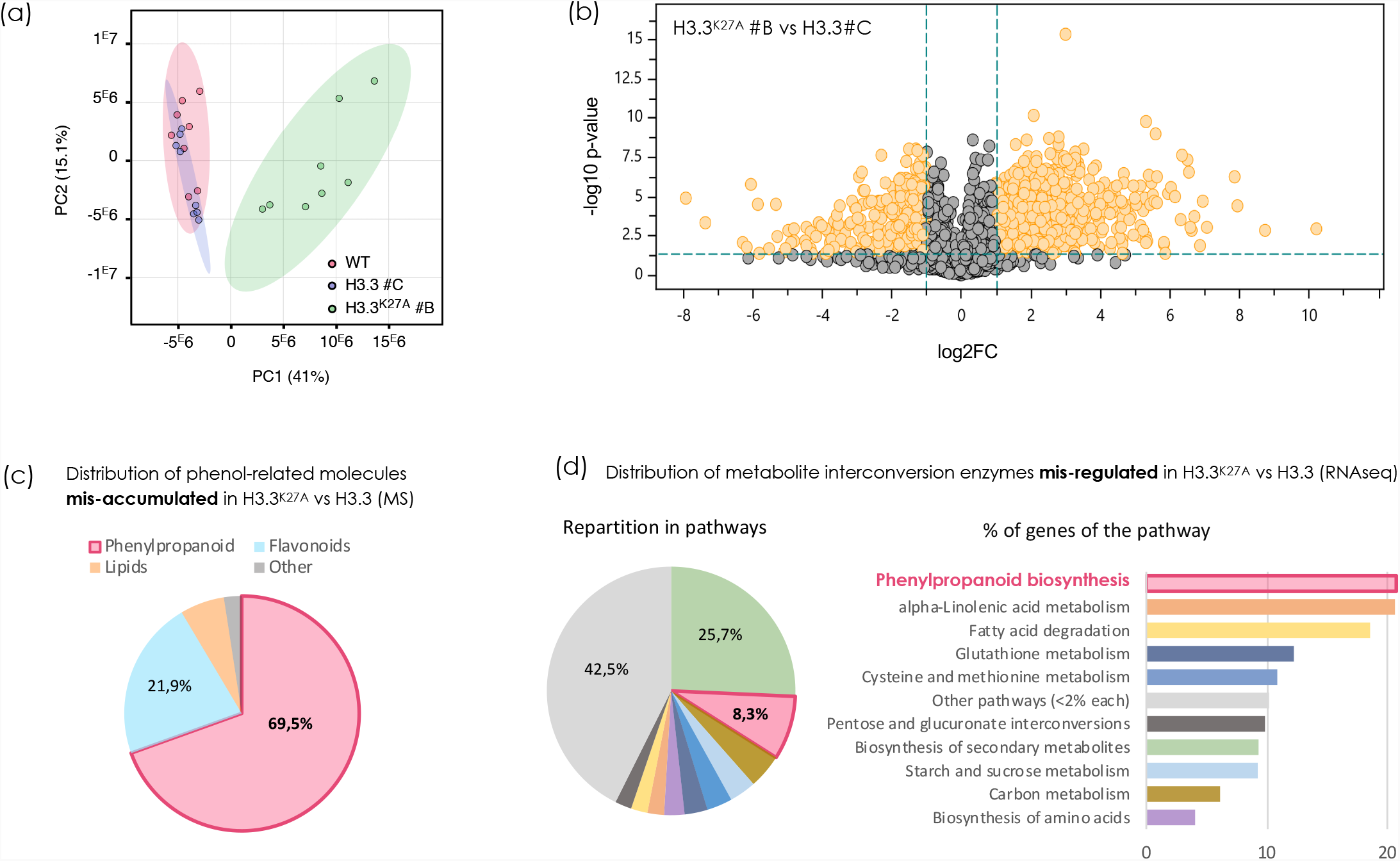
Metabolomics analyses on H3.3^K27A^ lines and correlation with the transcriptomics for interconversion enzymes. **(a)** 2-D score plot for principal component analysis (PCA) and **(b)** Volcano plot showing the differential enrichment of metabolites between WT, H3.3 #C and H3.3^K27A^ #B. Each point of the PCA represents a metabolite profile of an individual biological replicate. The PCA score plot was constructed using MetaboAnalyst 4.0. Data were analysed by range scalin. Each point in the volcano plot represents a metabolite (or bucket). Significant buckets were calculated with a fold change (FC) threshold of +/-2 and a p-value < 0.05 (indicated in orange). The distinct separation of H3.3^K27A^ #B in both (**a**) and (**b**) indicates the significant enrichment/depletion of more metabolites than in H3.3 #C compared to WT. **(c)** Pie chart illustrating the distribution of phenol-related molecules identified from our metabolomic data to be mis-accumulated in H3.3^K27A^ as compared to H3.3 and mapped to the KEGG compounds database. **(d)** Pie and bar charts illustrating the distribution of the 604 genes encoding for metabolite interconversion enzymes that were identified from our RNA-seq data to be mis-regulated in H3.3^K27A^ as compared to H3.3.

**Figure 5:**
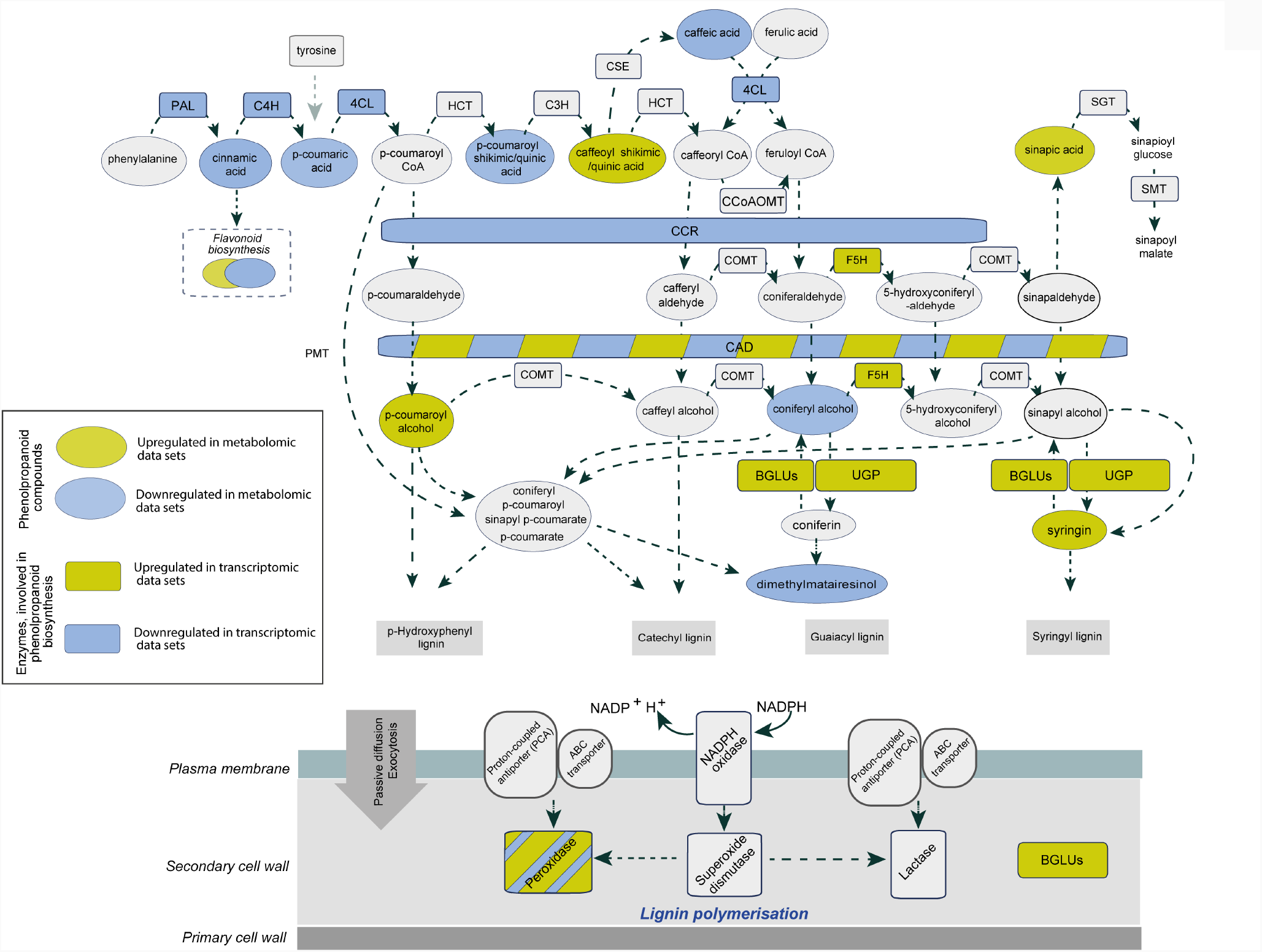
Phenylpropanoid biosynthesis pathway with transcriptome and metabolome for H3.3^K27A^ vs H3.3. Representation of transcriptomics and metabolomics data for phenylpropanoid biosynthesis (H3.3^K27A^ vs H3.3). Schematic illustration, depicting the elements of the Phenylpropanoid biosynthesis pathways that are detected to be mis-regulated in the metabolomic (ovals) and transcriptomic (rectangles) data sets, mapped to KEGG pathways database, with the colour code of mustard shade for upregulated and blue for downregulated elements. Abbreviations: PAL - phenylalanine ammonia-lyase; C4H - cinnamic acid 4-hydroxylase; 4CL - 4-coumarate-CoA ligase; HCT - Hydroxycinnamoyl-CoA shikimate/quinate hydroxycinnamoyl transferase; C3H - p-coumaroyl shikimate 3’ hydroxylase; CSE - caffeoyl shikimate esterase; CcoAOMT - caffeoyl CoA 3-O-methyltransferase; CCR - cinnamoyl-CoA reductase; COMT, caffeate/5-hydroxyferulate 3-O-methyltransferase; F5H - ferulate 5-hydroxylase; CAD - cinnamyl alcohol dehydrogenase; SGT - sinapato UDP-glucose sinapoyltransferase; SMT - 1-O- sinapoyl-β-glucose:L-malate O-sinapoyltransferase; BGLU -a-glucosidases; UGT - UDP-glycosyltransferase. Combined analysis of both data sets indicates the multiple levels of this pathway are affected in the H3.3^K27A^ plants.

### Metabolomics and transcriptomics highlight a function for H3.3K27 in phenylpropanoid biosynthesis

Because the largest proportion of genes mis-regulated in H3.3^K27A^ encode for metabolic interconversion enzymes, we launched an *in situ* metabolomic investigation of the H3.3^K27A^ lines. We determined the metabolome of WT, H3.3 (#C) and H3.3^K27A^ (#B) lines through a non-targeted approach in which buckets made of a mass/charge ratio and a retention time were used to describe molecules of interest. First, a Principal Component Analysis (PCA) indicated a clear separate clustering for H3.3^K27A^, while WT and H3.3 overlapped (Fig. 4a; Fig. S16a). In agreement with the PCA data, significantly more metabolites were found differently enriched in H3.3^K27A^ than in H3.3, compared to WT (i.e., fold change > +/- 2 and p-value < 0.05) (Fig. 4b and Fig. S16).

In depth analysis of mis-accumulated phenol-related molecules revealed a great enrichment in phenylpropanoids, which are core components of lignin (Fig. 4c, Tables S5 and S6). Interestingly, when returning to our RNA-seq data and detailing the pathways for metabolite interconversion enzyme-coding genes, we found a significant enrichment of genes involved in phenylpropanoid biosynthesis (Fig. 4d, Table S2). Strikingly, over 20% of genes of this GO pathway are mis-regulated (up-or down-regulated) in H3.3^K27A^. More precisely, we detected a decreased expression for several genes encoding enzymes of the lignin biosynthesis pathway, such as phenylalanine ammonia-lyase (PAL1, AT2G37040, log2FC*=*-1.67), cinnamic acid 4-hydroxylase (C4H, AT2G30490, log2FC*=*-1.16); 4-coumarate-CoA ligase (4CL, AT1G51680 log2FC=*-*1.05) (Table S2). This correlates positively with the lower content of the compounds produced by the corresponding enzymatic reactions (cinnamic and coumaric acids and their derivatives), observed in H3.3^K27A^. Also, the expression of multiple β-glucosidases (*BGLUs) and* UDP-glycosyltransferases (UGTs) is affected in H3.3^K27A^ (Table S2). Interestingly, BGLU45 and BGLU46 (log2FC=-1.33, -2.18 respectively), were previously shown to be involved in the hydrolysis of monolignol glucosides (the storage form of monolignols in Arabidopsis) (Chapelle *et al*., 2012) (Escamilla-Treviño *et al*., 2006).

In summary, we found a correlation between misrepresented metabolic compounds, corresponding mis-regulated genes and the xylem over-proliferation in stems of the H3.3^K27A^ lines.

## Discussion

### The H3.3^K27A^ lines reveal developmental and molecular functions for H3.3K27

The substitution of K27 into an Alanine residue in the H3.3 protein proved to be highly instrumental in discovering novel functions for the histone Lysine residue in plants. It illustrates how loss of histone modifications, independently from loss of histone-modifying enzymes, can impact biological processes such as gene expression and biosynthetic pathways. While the H3.3^K27A^ lines displayed developmental defects reminiscent of PRC2 loss-of-function, such as early flowering, leaf morphology defects and shorter stems, our approach revealed novel developmental features as well. Indeed, we found that the H3.3^K27A^ lines are not affected in flower number or morphogenesis, but have apparent defects in cell morphology and elongation in the stem epidermis, combined with a modified cell layer patterning within the stem, and faster calli proliferation.

Our cytological analyses did not reveal a profound impact of H3.3^K27A^ on hetero/euchromatin spatial distribution, but our molecular analyses unveiled changes in the level of euchromatin marks in the nucleus. Indeed, while the H3.3^K27A^ variant led to a decreased loading of the H3K27me3 and H3K27ac euchromatic histone marks, it did not impact the level of the constitutive heterochromatin H3K27me1 mark, likely correlating with the globally maintained nuclear and chromatin organization in the H3.3^K27A^ lines. In *Arabidopsis thaliana*, the preferred substrate of the H3K27 monomethylases ATXR5 and ATRX6 (Jacob *et al*., 2009), which are unique to plants, was reported to be the H3.1 variant (Bieluszewski *et al*., 2021). *In vitro* work showed that the preference of ATRX5/6 for H3.1 over H3.3 is due to a single amino acid difference at position 31 of H3, with Alanine promoting H3.1 methylation, while Threonine in H3.3 blocks methylation (Jacob *et al*., 2014). Here we thus provide an *in vivo* evidence for the preferential K27 mono-methylation of H3.1 versus H3.3, indicating an efficient H3 variant sorting for the deposition of this mark.

H3.1 and H3.3 variants differ in only four amino acids, yet they have distinct deposition patterns within the chromatin, where they exert different functions (Shi *et al*., 2011) (Probst *et al*., 2020). While H3.1, which is rather enriched at silent regions and predominantly expressed and incorporated during the S-phase, H3.3 is expressed and deposited throughout the cell cycle, independently from DNA replication (Ahmad & Henikoff, 2002)(Jiang & Berger, 2017a).

Interestingly, our data also indicates that in Arabidopsis, the overexpression of canonical histones such as H3.3 has no effect on the plant growth and development (i.e. the H3.3 control lines have a wild-type phenotype), while this is not the case in yeast (Singh *et al*., 2010). Another interesting point is the robustness of the observed phenotypes, for their types and strength, over generations and between individuals of a same line. This high penetrance indicates that either, H3.3^K27A^ and H3.3 histones are consistently loaded at individual chromatin positions, and/or that the regulation of some genes and corresponding pathways is more sensitive than others to the H3.3^K27A^ mutated histone.

The H3K27me3 mark has been extensively reported as contributing to the maintenance of epigenetic states for expression program memory over cell cycles in animals and plants (Mozgova *et al*., 2015)(Hugues *et al*., 2020), rather than initiating gene expression changes (Lafos *et al*., 2011)(Engelhorn *et al*., 2017). The developmental phenotypes observed in H3.3^K27A^ lines, including early flowering and stem cell fate defects, likely reveals a role for H3.3K27 in establishing the regulation of target genes to be switched for a developmental transition, rather than in maintaining an already established gene expression state, hence corresponding cell fate. Overexpression of H3.3^K27A^ may mostly impact the initial deposition of repressive H3K27me3 for gene expression reprogramming, before replacement by the replication-coupled H3.1 for establishment of a maintained repressed state over cell divisions (Jiang & Berger, 2017b).

Mutating H3.3 therefore offers an experimental advantage for depicting the specific function of a histone variant, while ‘killing’ histone modifications in a variant-selective manner. As a matter of fact, H3.3^K27A^ variants, but not H3.1^K27A^ variants, induced striking developmental phenotypes in *Arabidopsis thaliana* plants (data not shown).

### H3.3K27 controls the capacity to regenerate tissues

Culturing H3.3^K27A^ leaves on CIM, which contains two hormones providing a strong signal to divide, resulted in rapid callus formation with the outcome of larger calli than the controls. Enhanced callus formation was also observed in plants over-expressing the atypical H3.15 variant lacking Lysines at positions 4 (N4) and 27 (H27) (Yan *et al*., 2020). Authors speculated that up-regulation of *WOX11* and *LATERAL ORGAN BOUNDARIES DOMAIN18* (*LBD18*) genes promoted the callus formation. Two of the four LBD TFs (LBD16 to 18, and LBD29) that were identified as enhancers of callus formation (Fan *et al*., 2012) are marked by H3K27me3 in WT callus and are up-regulated in H3.3^K27A^. Previously, the up-regulation of *LBD29* in *myb94myb96* double mutant was shown to stimulate cell proliferation on CIM (Dai *et al*., 2020). Therefore, *LBD29* up-regulation in H3.3^K27A^ (log2FC=7.4) may also explain the highly proliferating callus phenotype. Moreover, the bZIP59–LBD complex downstream gene *FAD-binding Berberine* (*FAD-BD*), which contributes to callus formation (Xu *et al*., 2018), is also up-regulated in H3.3^K27A^ (log2FC=3.8). Altogether the callus phenotype of H3.3^K27A^ highlights the role of H3.3 K27 in controlling cell proliferation on CIM.

Our H3.3^K27A^ lines reveal another, newly discovered, cell fate property: the failure to regenerate shoots from calli transferred to SIM. Indeed, H3.3^K27A^ calli grown on SIM regenerate structures reminiscent of *clfswn* double mutants (Chanvivattana *et al*., 2004), thus establishing a link between the impaired regenerative capacity of H3.3^K27A^ and the decrease in H3K27me3. The acquisition of new fate upon regenerative stimulus requires the establishment of specific gene expression patterns, which involves mechanisms to silence the previous cell identity genes and reinforce lineage commitment (Eshed Williams, 2021). In the H3.3^K27A^ up-regulated genes, we identified many developmental TFs from different pathways, some of them being sufficient for direct differentiation and organogenesis.

Among these, are members of the MIKC_MADS or YABBY TF families, which corresponding genes are marked by H3K27me3 in WT callus. We thus propose that the failure of H3.3^K27A^ calli to regenerate shoot is due to a lack of mechanisms that silence these TFs to allow the shoot programme to dominate.

### H3.3K27 is a critical epigenetic checkpoint for stem morphogenesis and lignin biosynthesis

With an integrative approach gathering transcriptomics and metabolomics, we found a correlation between mis-regulated genes and mis-represented metabolites, in connection with some of the phenotypes observed in H3.3^K27A^. Indeed, short-stemmed H3.3^K27A^ plants displayed morphologic defects both at the epidermis and the inner stem cell layers. In particular, histological analyses on stem longitudinal and cross sections revealed thickening of the lignified tissues layers and vasculature bundles, indicating mis-regulation of cell specification. Interestingly, we found that the expression of *WOX4* is reduced in the stems of H3.3^K27A^ plants. WOX4 is known to initiate the self-division of cambium cells, thereby controlling vascular cell proliferation and secondary growth of the stem both in Arabidopsis and Poplar (Hirakawa *et al*., 2010) (Kucukoglu *et al*., 2017). A decrease in *WOX4* expression is known to promote xylem and phloem differentiation, which is what we observe in the stems of H3.3^K27A^ lines. Strikingly, out of the 266 genes identified as downregulated in the inflorescence stems of *wox4* (Suer *et al*., 2011), 55 genes are also mis-regulated in H3.3^K27A^ plants (33 of them are upregulated and 22 are downregulated in H3.3^K27A^), which represents a significant overlap (RF= 1.8, p-val. < 6.696e-06). Among the genes downregulated in *wox4* and upregulated in H3.3^K27A^, 24% are covered with H3K27me3 (8 of 33), compared to 9% (2 of 22) for the commonly downregulated genes (Table S7). This once again supports an effect of the H3.3^K27A^ variant on H3K27 tri-methylation. Further supporting the links with the observed plant phenotypes, 41% (9 of 22) of the genes downregulated in both H3.3^K27A^ and *wox4*, are known to be expressed in xylem.

Finally, the fact that many genes involved in multiple biosynthetic pathways were found mis-regulated in H3.3^K27A^ prompted us to explore its metabolome. Metabolomics is a powerful approach to explore how genetic diversity may affect phenotypic variations in plants, but was until now never used in the context of chromatin. It revealed a mis-accumulation of phenol-related molecules, among which phenylpropanoid components were the most represented, thus confirming the effect of the identified transcriptomic mis-regulation on the production of plant metabolic components.

### Transcriptomics data reveal overlap and differences between H3.3^K27A^ and emf2 PRC2 mutant lines

The role of H3.3K27 is revealed by the mis-regulation of more than 3000 genes, out of which more than 1700 are up-regulated, and 620 (>36%) are direct targets of H3K27me3 or of PRC2. In general, studies on PRC2 mutants such as *clf, swn* or *emf2* do not show such a good overlap even when considering PRC2-bound genes (Shu *et al*., 2019) (Mandel *et al*., 2022). Comparing our RNA-seq data with dataset obtained on *emf2* calli, we found that there are many more up-regulated genes in the H3.3^K27A^ calli than in the PRC2 loss-of-function mutant. Among the TF families up-regulated in H3.3^K27A^, whose proportion increases when looking at genes covered with H3K27me3 and up-regulated in *emf2*, the fraction of MADS-encoding genes increases the most. This is in agreement with the many reports showing that MADS TFs are key targets for PRC2 regulation (Zhang *et al*., 2018)(Kim *et al*., 2012)(Engelhorn *et al*., 2017). Interestingly and differently to the TF-encoding genes up regulated in H3.3^K27A^, most of the down-regulated TF-encoding genes also known to be H3K27me3 targets, encode for HTH and bHLH families only.

Our approach using the non-methylable H3.3^K27A^ variant thus reveals the mis-expression of a larger set of H3K27me3-regulated genes than those that study transcriptomics of PRC2 mutants. The H3.3^K27A^-encoding gene is driven by a strong constitutive promoter known to conduct gene expression in all cell types of all the plant organs, in a comparable manner. Therefore, we assume that the histone variant is equally expressed in all the tissues analysed in our study and has been integrated into the chromatin in a random fashion, consistently in all cell types. Consequently, our approach provides an advantage on the use of mutants in mark writers that are subjected to specific tissue or can vary in expression level between different cell types (de Lucas *et al*., 2016).

In brief, our study reveals that the histone H3.3 K-to-A mutation at K27 acts as a dominant-negative gain-of-function mutation in plants. Expression of the H3.3^K27A^ variant leads to unique developmental aberrations, part of them reminiscent of PRC2 mutant phenotypes and part of them novel. They include early flowering, leaf morphology defects, impaired stem elongation and cell layer patterning. Transcriptomic and metabolomic analyses allowed us to correlate newly discovered phenotypes to defective biosynthesis pathways, in particular that of the phenylpropanoid pathway related to lignin biosynthesis. Given the highly conserved Histone 3 sequence across the plant (and animal) kingdom(s), mutations at K27 provide a mean to alter epigenetic landscapes in organisms where histone methyltransferases are uncharacterized or where genetic studies are challenging due to multiple redundancies and multifaceted functions.

## Acknowledgements

We thank Muriel Kabus, Pauline Brun, Elda Bauda for help with plant culture and selection of transgenic lines, and Emmanuel Thévenon for synthetic DNA cloning and plasmid amplification. This work was supported by the Agence Nationale de la Recherche (ANR-18-CE20-0011-01, PRC project REWIRE to CCC, AB and MEC), the Grenoble Alliance for Cell and Structural Biology (ANR-10-LABX-49-01) and the Giant International Internship program.

## Author Contribution

Design of the research strategy: CCC and LEW. Research experiments: KF (flowering time, organogenesis, apex and stem measurements, histology, RT-qPCR for stem features), MLM (genetics, genotyping and RT-PCR for selection of transgenic lines), DT (root cytological work), AB (flowering time, western blots, measurements and RT-qPCR for flowering time), EP (selection and primary analyses of transgenic lines), NI (stem SEM), NLM (callus and regeneration), CV (metabolomics), CCC (construction, selection and phenotypic analyses of transgenic lines, plant imaging, western blot). Data collection and analysis: KF (statistical analyses of phenotypes, image processing and analysis, RNA-seq analysis, metabolomics analysis), AB (RT-qPCR, RNA-seq analysis, metabolomic analysis), AF (RNA-seq collection and analysis), CV (metabolomic collection and analysis), LEW and CCC (population overlaps, enrichments and GO analyses on RNA-seq and metabolomics). Data interpretation: KF (flowering, organogenesis, stem, RT-qPCR, reconstruction of the phenylpropanoid biosynthesis pathways from transcriptomics and metabolomics), AB (flowering, RT-qPCR, RNA-seq, metabolomics), LEW (callus, stem SEM, RNA-seq) and CCC (flowering, RT-qPCR, RNA-seq, metabolomics). Funding for the study: CCC, AB, MEC and LEW. CCC wrote the manuscript with the help of LEW, AB and KF. All authors read and approved the final manuscript.

## Data Availability

Transcriptomics and metabolomics data that support the findings of this study are available as article supplementary information material (Supplementary Tables). Raw data are available from the corresponding author upon reasonable request (e.g. from editor and/or reviewers), and will be freely accessible after deposition on a repository platform upon paper acceptance.

