## Supplementary material for "Lysine 27 of histone H3.3 is a fine modulator of developmental gene expression and stands as an epigenetic checkpoint for lignin biosynthesis in Arabidopsis": Suppl Figures

Supplementary Figure S1

### Sequences of the H3.3 and H3.3$^{K27A}$ – encoding Transgenes

**>H3.3 (control)**

ATGGCTCGTACTAAGCAAACAGCTCGTAAGTCTACTGGAGGAAAGGCTCCTAGGAAGCAGCTTGCTACAAA

Ggtaagactcgggctctcacatgtgatctgagtagcttgataaacacatttctagatttgttctaattggt

ggatgttttaatttaagGCTGCACGTAAGTCTGCACCAACCACTGGAGGAGTCAAGAAGCCCCATCGTTAC

CGTCCAGGAACTGTTGCACTACGgtatgcaatctgtttcttccccaaattcaattgtgtttttactttatt

gatatcgtcattgattaaactctgtttctctgttttattgatcttgattgacagTGAAATTCGTAAGTACC

AGAAGAGTACCGAGTTGCTGATCAGGAAGCTCCCTTTCCAGAGGCTAGTTCGTGAGATTGCCCAGGATTTC

AAGgtaatgtgtctgttcgtctgattcagtctacaaatggatttaactttttacttgttttgtcataacct

tgtgttcttatcttgtctctttgattgcagACTGACTTGCGTTTCCAGAGCCATGCTGTGCTTGCACTCCA
oMLMc2 →

GGAGGCTGCTGAAGCATACCTTGTGGGTCTCTTTGAGGACACTAACCTCTGCGCCATCCATGCCAAGCGTG

TGACCATAATGCCCAAAGACATTCAACTTGCACGTAGAATTAGGGGTGAACGTGCTTAA
← oMLMc8

**>H3.3$^{K27A}$ (variant)**

ATGGCTCGTACTAAGCAAACAGCTCGTAAGTCTACTGGAGGAAAGGCTCCTAGGAAGCAGCTTGCTACAAA

Ggtaagactcgggctctcacatgtgatctgagtagcttgataaacacatttctagatttgttctaattggt

ggatgttttaatttaagGCTGCACGTGCTTCTGCACCAACCACTGGAGGAGTCAAGAAGCCCCATCGTTAC

CGTCCAGGAACTGTTGCACTACGgtatgcaatctgtttcttccccaaattcaattgtgtttttactttatt

gatatcgtcattgattaaactctgtttctctgttttattgatcttgattgacagTGAAATTCGTAAGTACC

AGAAGAGTACCGAGTTGCTGATCAGGAAGCTCCCTTTCCAGAGGCTAGTTCGTGAGATTGCCCAGGATTTC

AAGgtaatgtgtctgttcgtctgattcagtctacaaatggatttaactttttacttgttttgtcataacct

tgtgttcttatcttgtctctttgattgcagACTGACTTGCGTTTCCAGAGCCATGCTGTGCTTGCACTCCA
oMLMc2 →

GGAGGCTGCTGAAGCATACCTTGTGGGTCTCTTTGAGGACACTAACCTCTGCGCCATCCATGCCAAGCGTG

TGACCATAATGCCCAAAGACATTCAACTTGCACGTAGAATTAGGGGTGAACGTGCTTAA
← oMLMc8

### Sequence of *HTR5* Endogene coding for H3.3

**>*HTR5* (At4g40040)**

ATGGCTCGTACTAAGCAAACAGCTCGTAAGTCTACTGGAGGAAAGGCTCCTAGGAAGCAGCTTGCTACAAA

Ggtaagactcgggctctcacatgtgatctgagtagcttgataaacacatttctagatttgttctaattggt

ggatgttttaatttaagGCTGCACGTAAGTCTGCACCAACCACTGGAGGAGTCAAGAAGCCCCATCGTTAC

CGTCCAGGAACTGTTGCACTACGgtatgcaatctgtttcttccccaaattcaattgtgtttttactttatt

gatatcgtcattgattaaactctgtttctctgttttattgatcttgattgacagTGAAATTCGTAAGTACC

AGAAGAGTACCGAGTTGCTGATCAGGAAGCTCCCTTTCCAGAGGCTAGTTCGTGAGATTGCCCAGGATTTC

AAGgtaatgtgtctgttcgtctgattcagtctacaaatggatttaactttttacttgttttgtcataacct

tgtgttcttatcttgtctctttgattgcagACTGACTTGCGTTTCCAGAGCCATGCTGTGCTTGCACTCCA
oMLMc2 →

GGAGGCTGCTGAAGCATACCTTGTGGGTCTCTTTGAGGACACTAACCTCTGCGCCATCCATGCCAAGCGTG

TGACCATAATGCCCAAAGACATTCAGCTCGCTCGCAGGATCAGAGGAGAACGTGCTTAA
← oMLMc11

Supplementary Figure S2

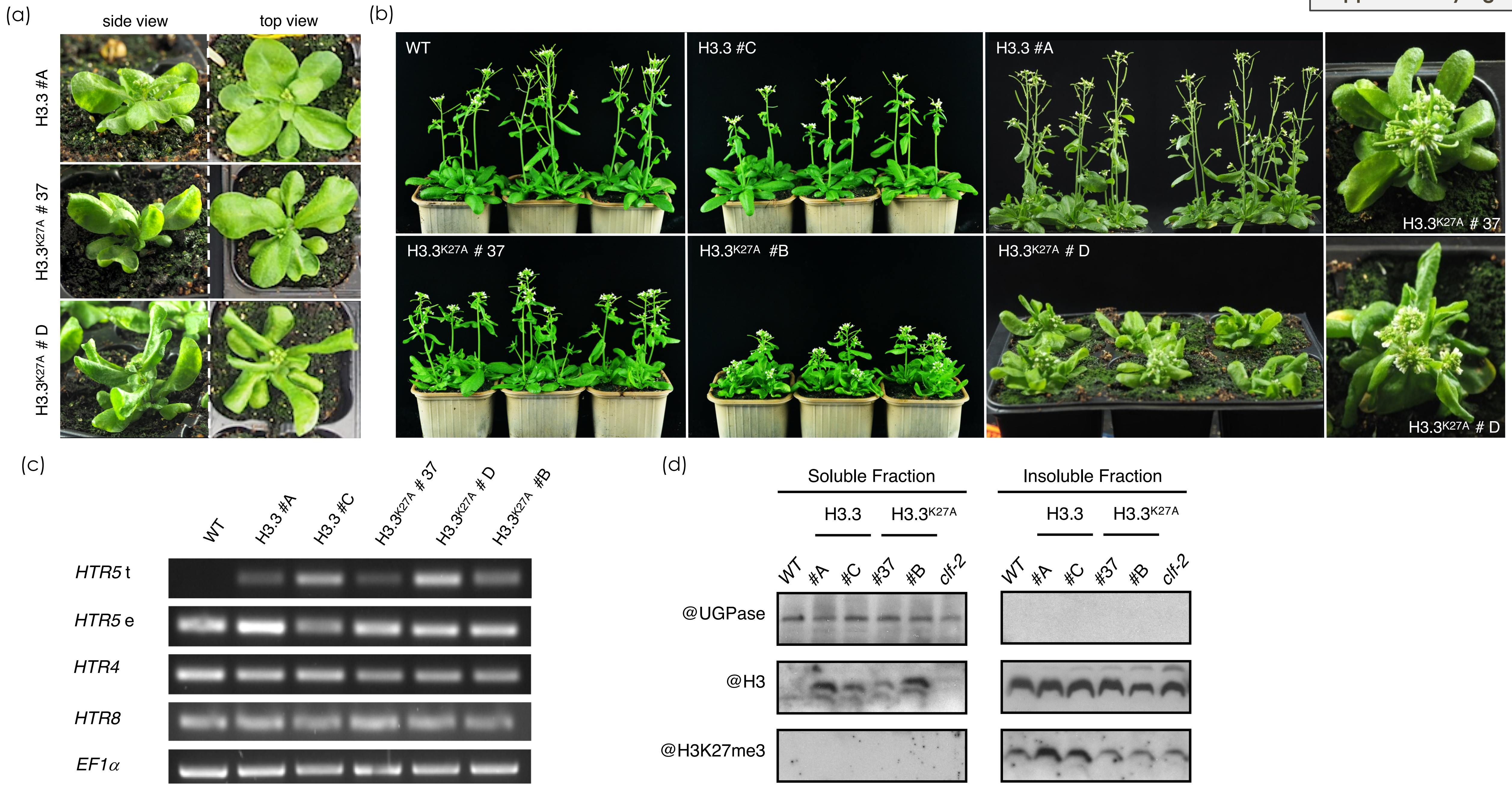

### Supplementary Figure S3

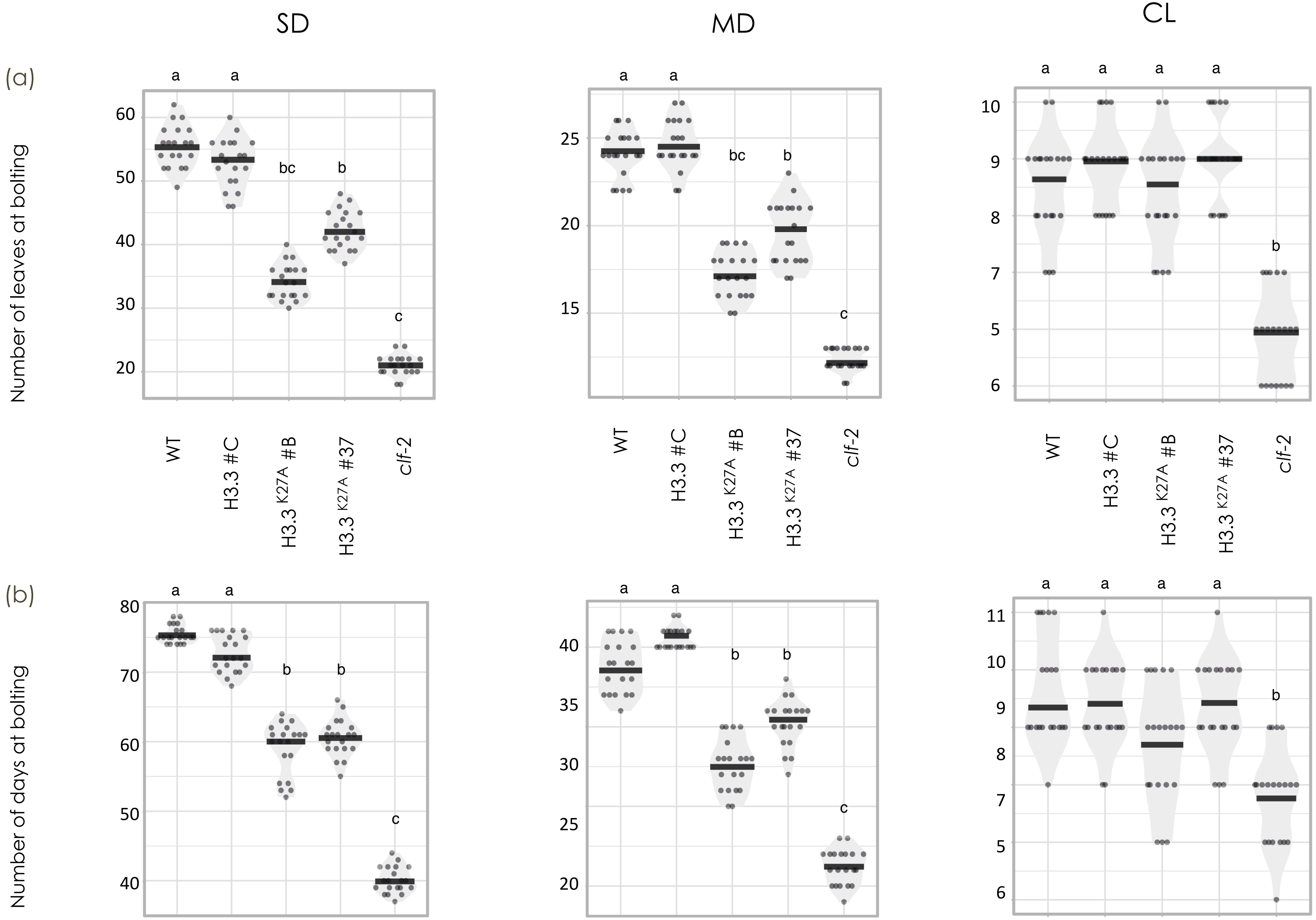

**Supplementary Figure S4**

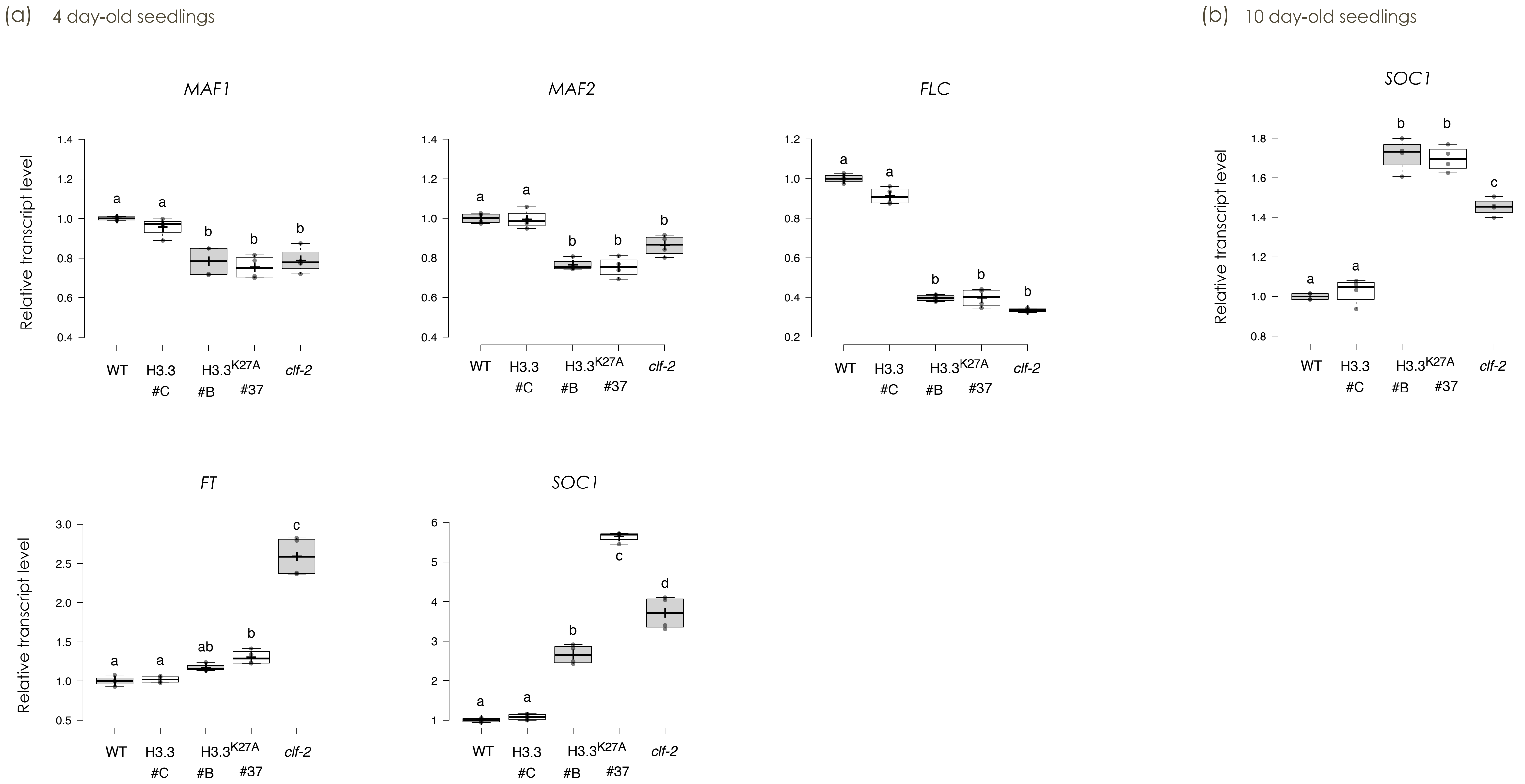

Supplementary Figure S5

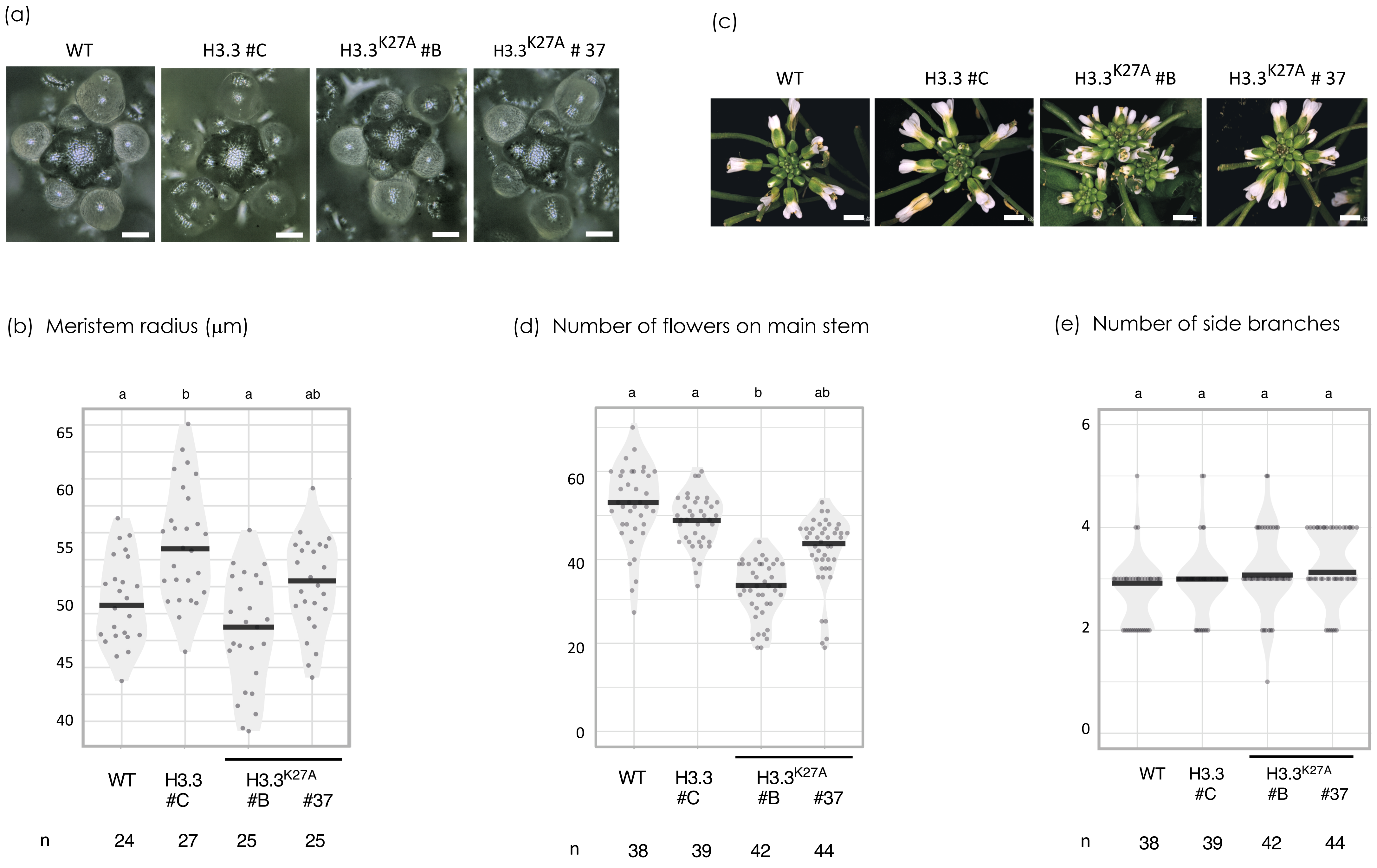

Supplementary Figure S6

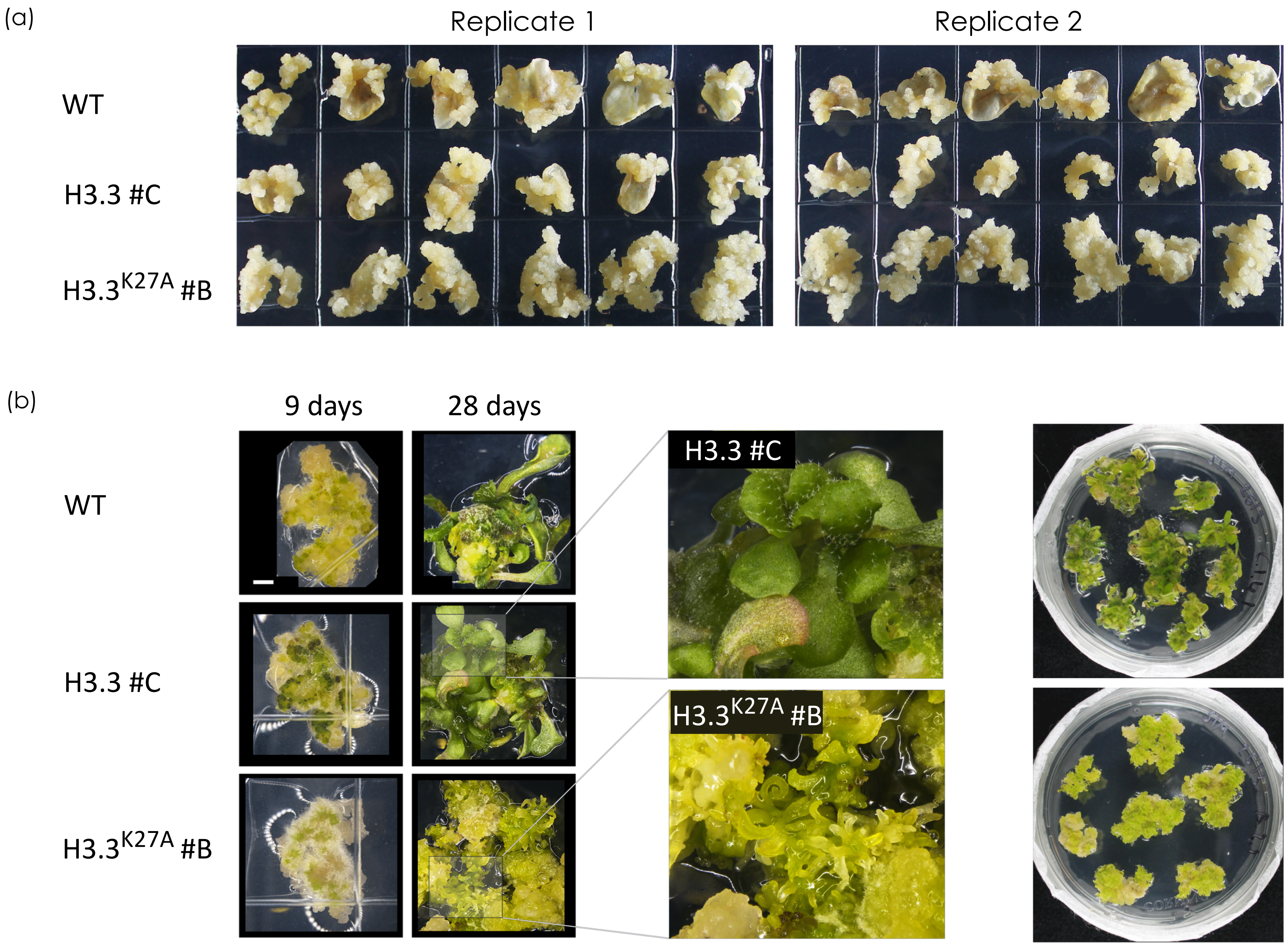

Supplementary Figure S7

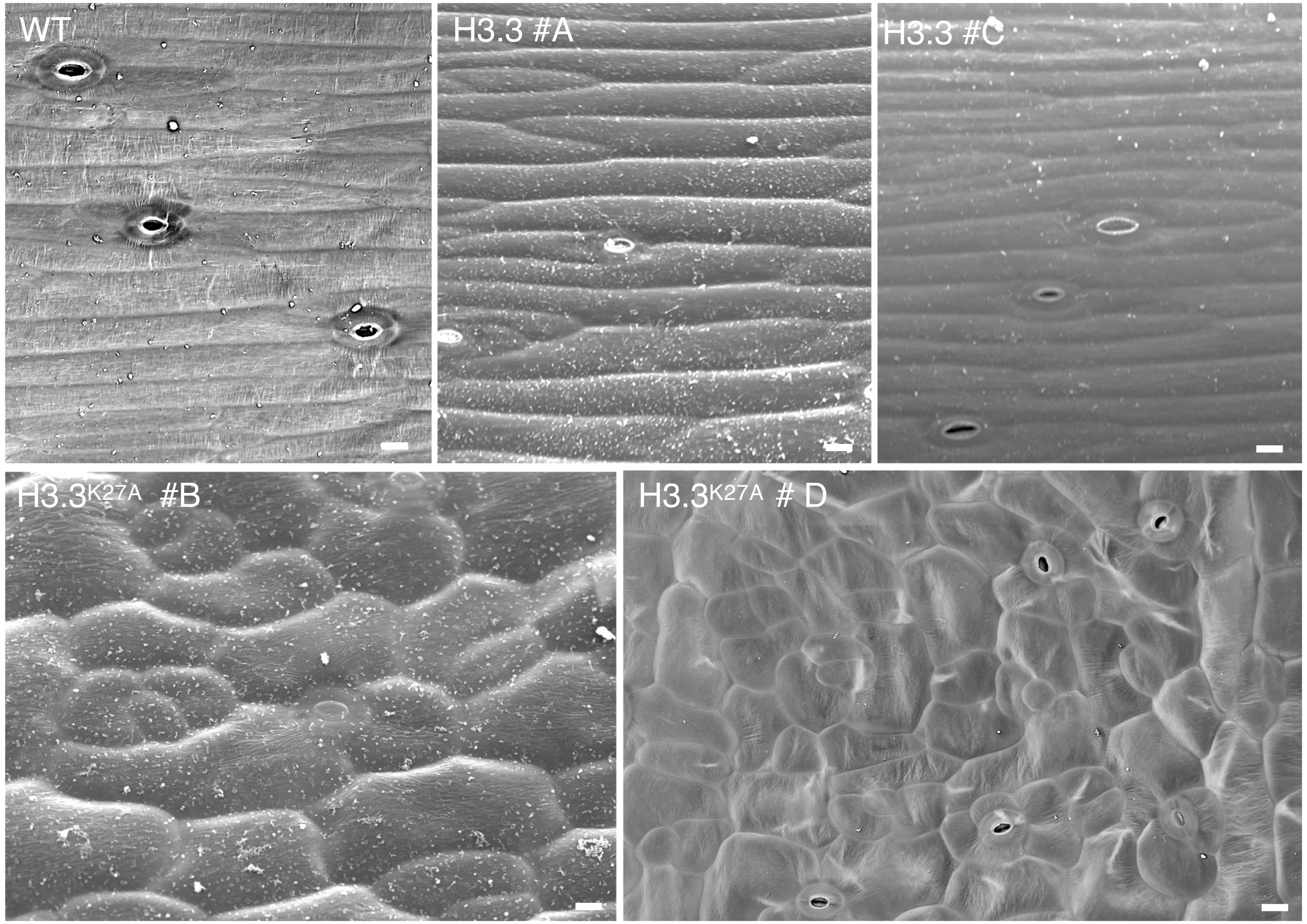

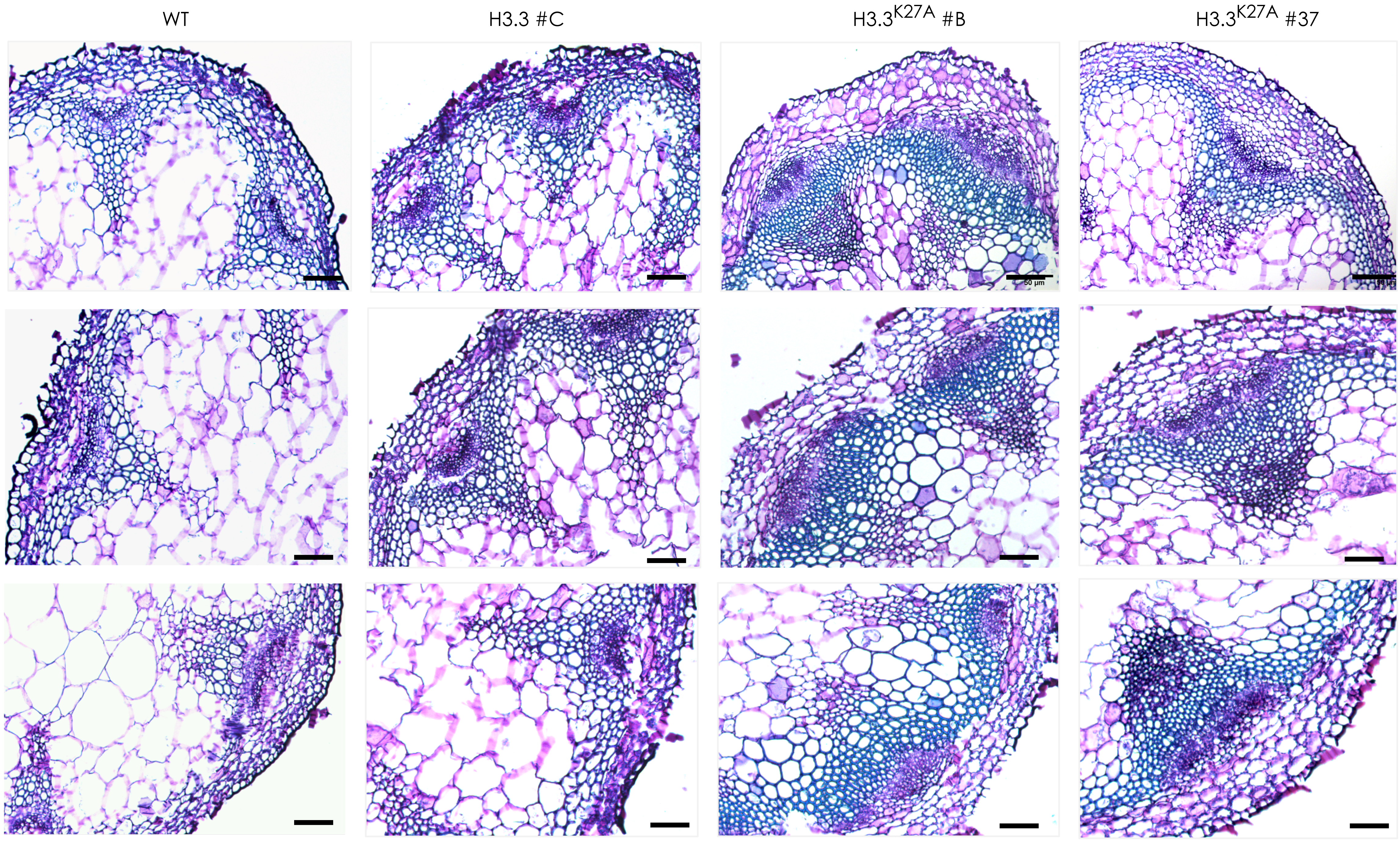


**Supplementary Figure S8**

Supplementary Figure S9

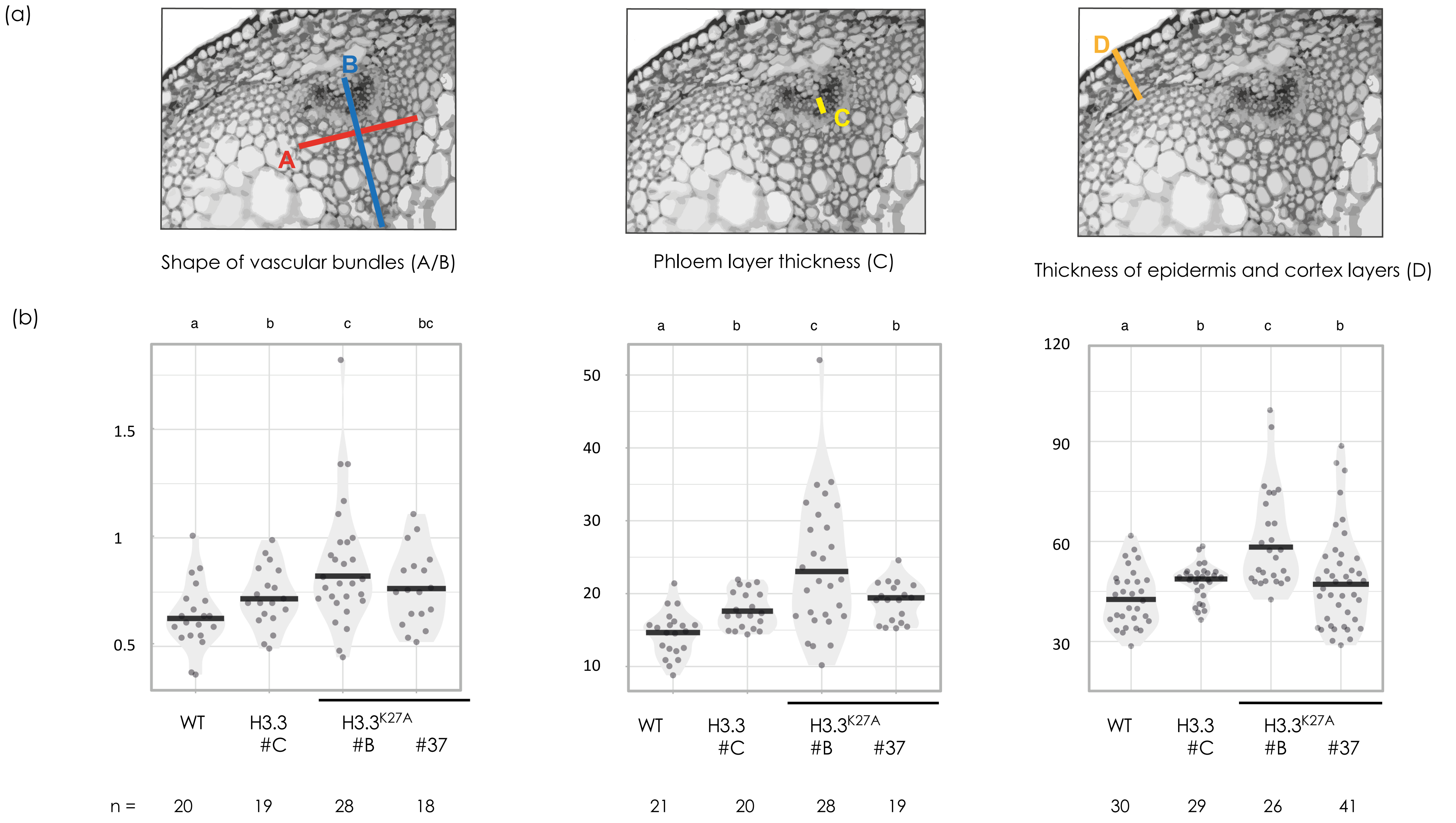

**Supplementary Figure S10**

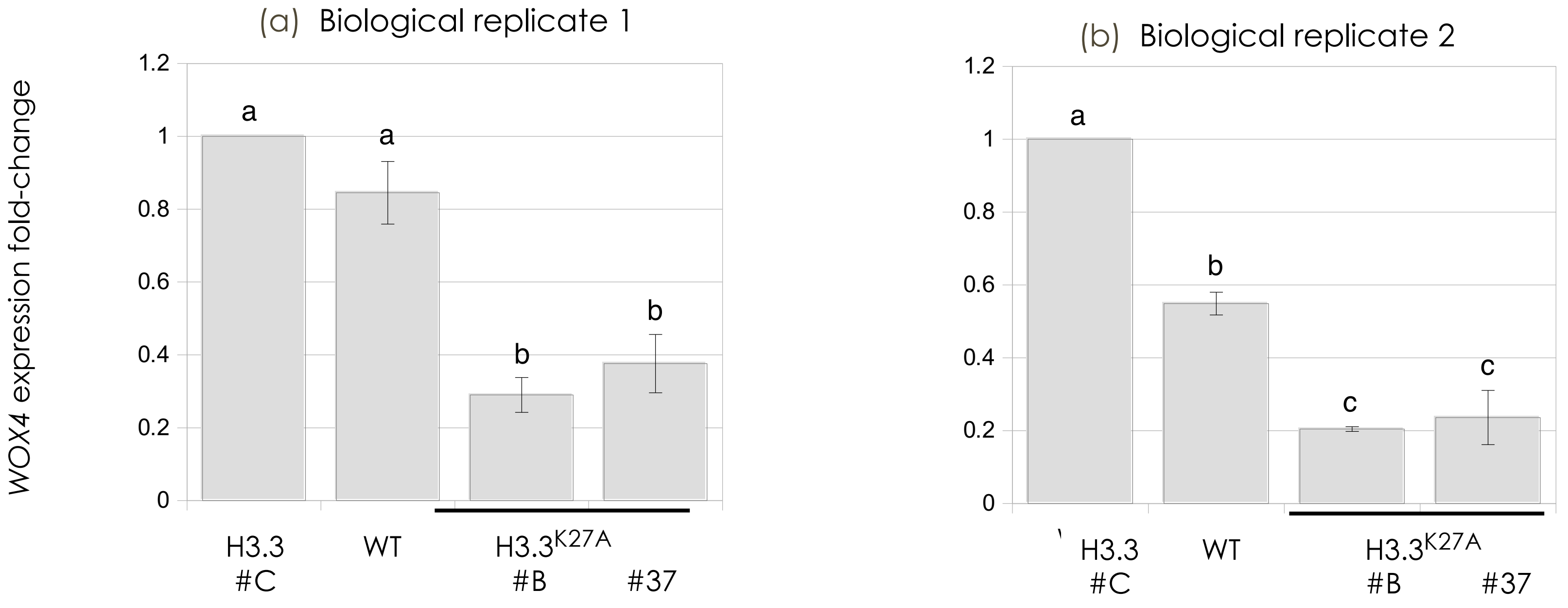

**Supplementary Figure S11**

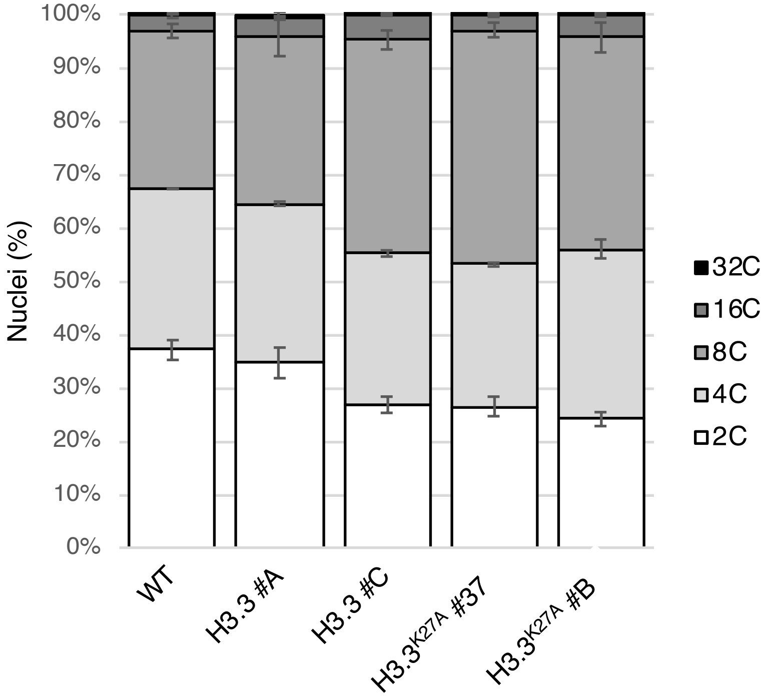

Supplementary Figure S12

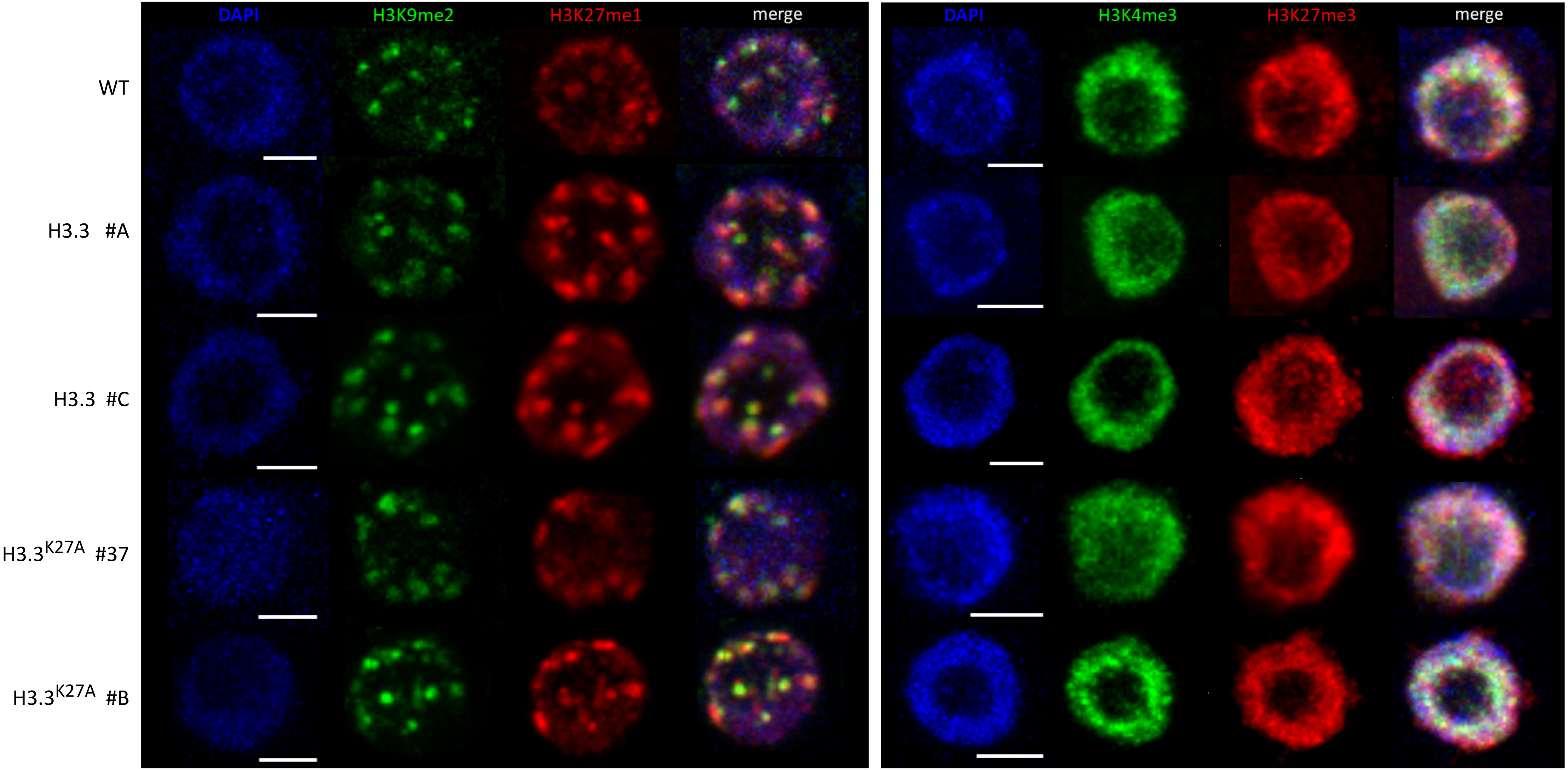

Supplementary Figure S13

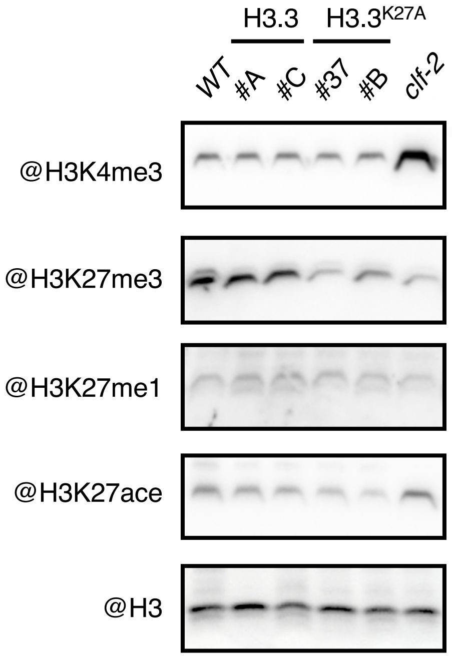

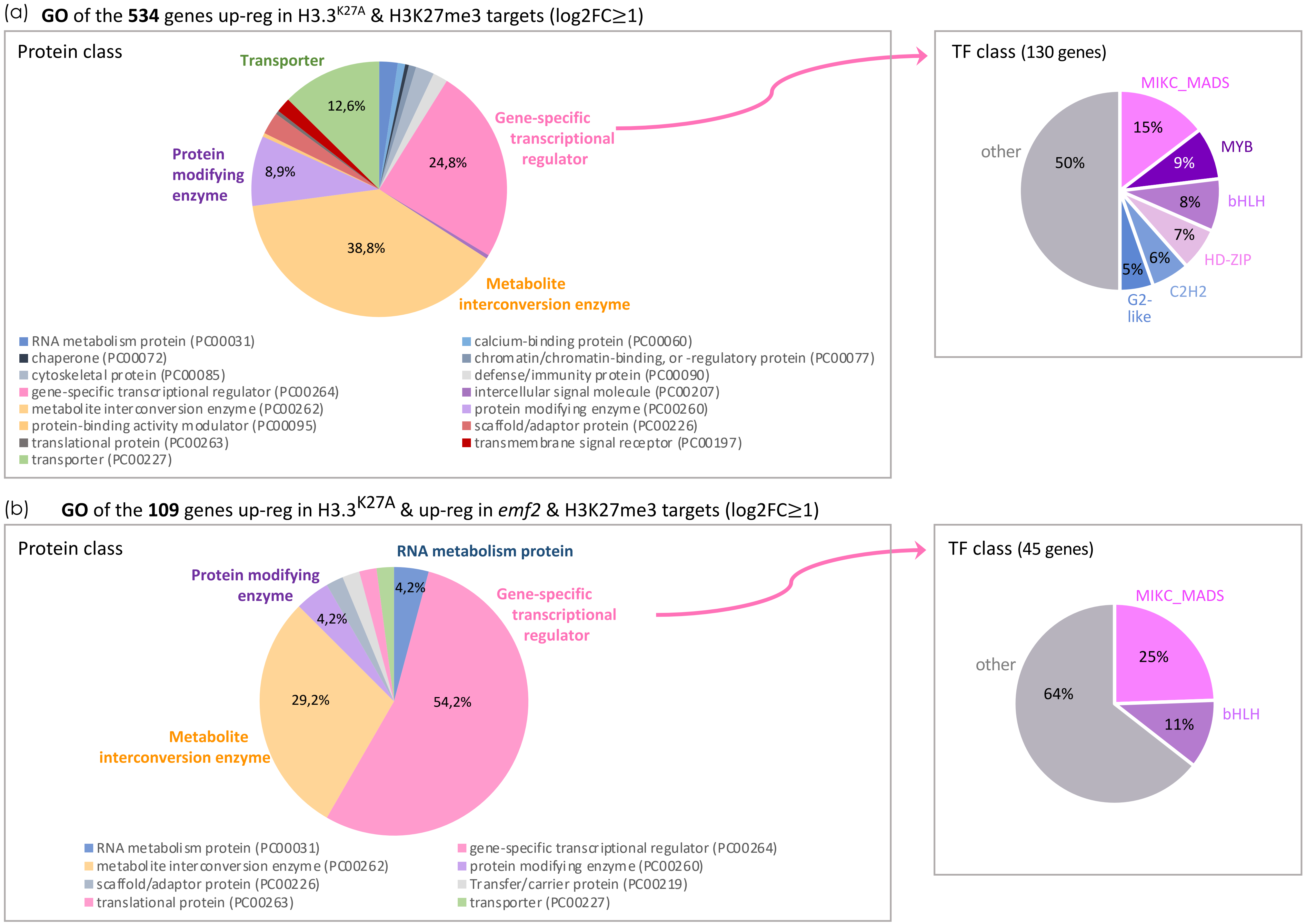


Supplementary Figure S14

**Supplementary Figure S15**

(a) **GO** of the **86** genes down-reg in H3.3$^{K27A}$ & H3K27me3 targets (log2FC ≤-1 )

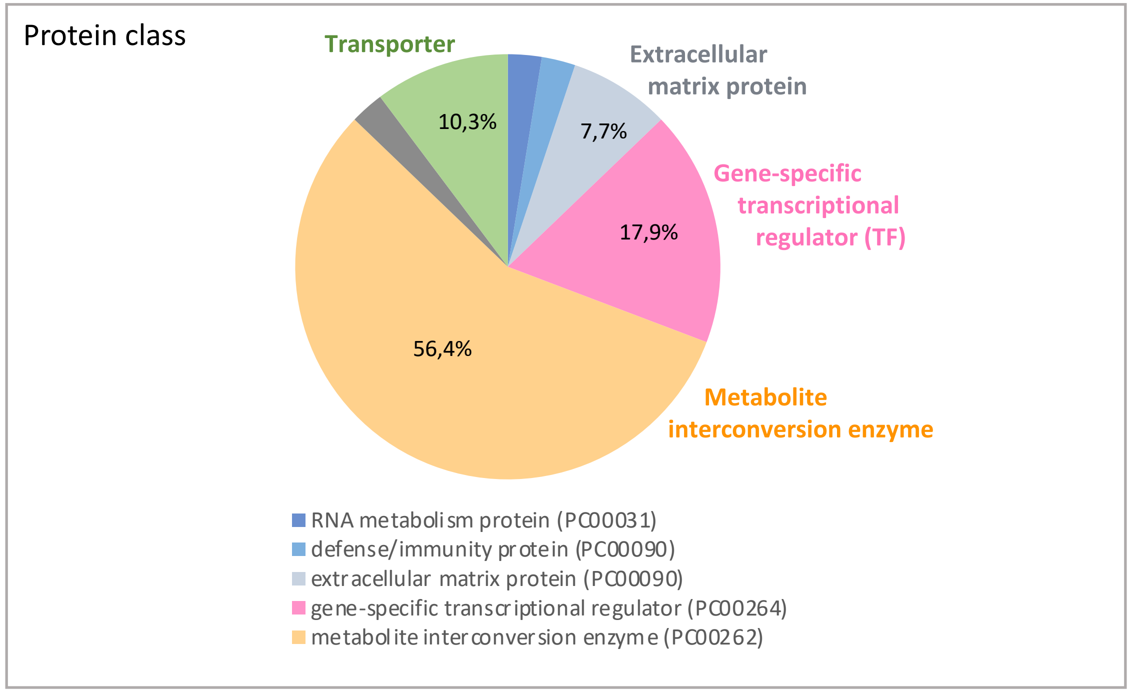


(b) **GO** of the **18** genes down-reg in H3.3$^{K27A}$ & down-reg in *emf2* & H3K27me3 targets (log2FC ≤-1 )

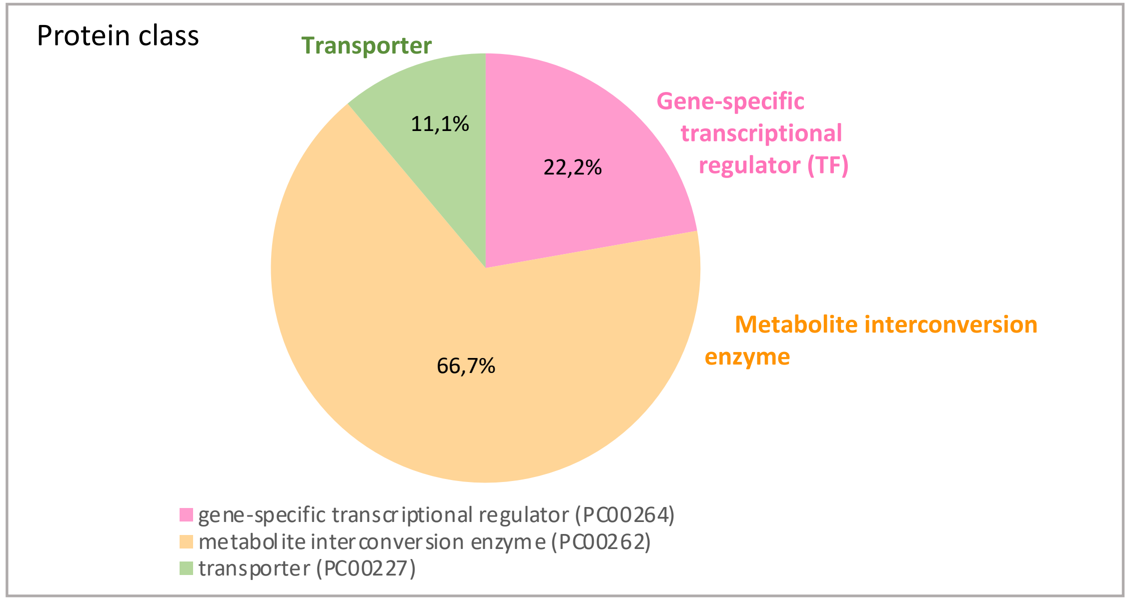


(c) Proportion of TFs (%) in the various populations of down-regulated genes in overlap with H3K27me3 targets and/or *emf2* down-regulated genes

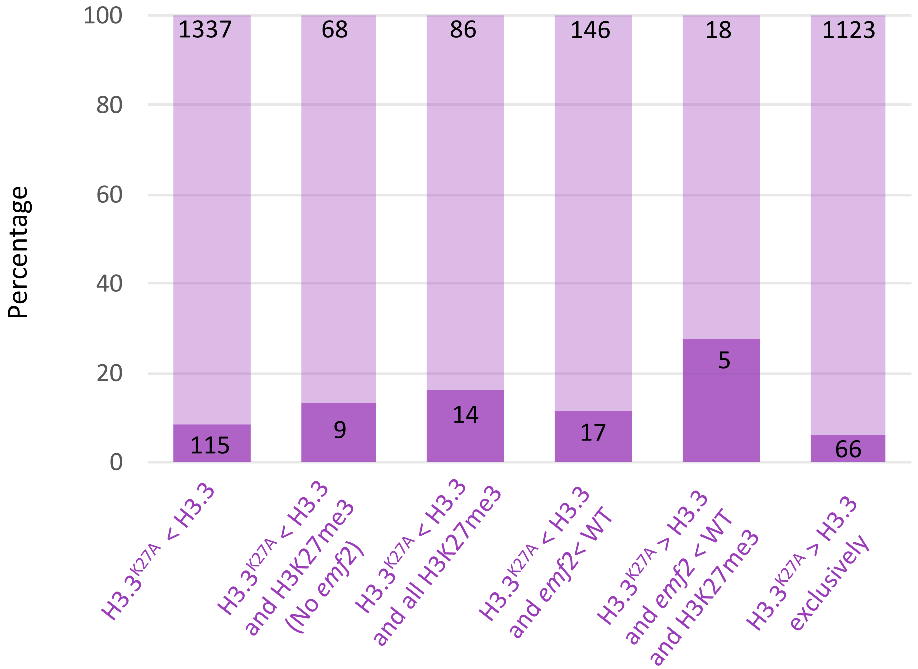

**Supplementary Figure S16**

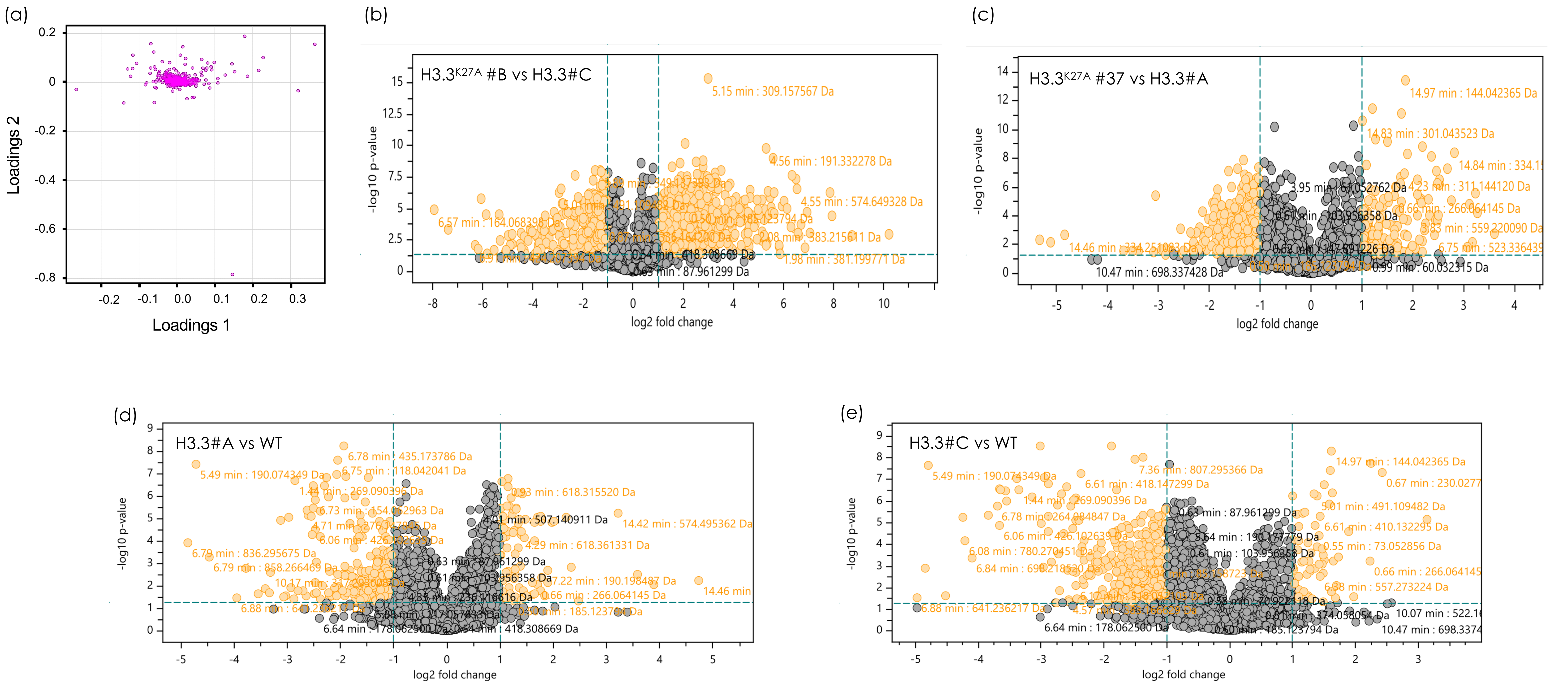
