## Supplementary material for "Lysine 27 of histone H3.3 is a fine modulator of developmental gene expression and stands as an epigenetic checkpoint for lignin biosynthesis in Arabidopsis": Suppl Information

*New Phytologist* Supporting Information

Article acceptance date: Click here to enter a date.

The following Supporting Information is available for this article:

**Fig. S1 Sequences of the H3.3- and H3.3$^{K27A}$ -encoding transgenes (designed from the *HTR5* sequence) and of the H3.3-encoding *HTR5* (At4g40040).**

**Fig. S2 Selection of sets of H3.3 and H3.3K27A lines with comparable transgene expression and analysis of H3 protein accumulation in the soluble vs insoluble fraction.**

**Fig. S3 H3.3$^{K27A}$ lines flower early in mid-day (MD) and long-day (LD) conditions.**

**Fig. S4 Expression analysis of key flowering genes in the H3.3 control (H3.3 #C) and H3.3K27A (H3.3K27A #B and #37) compare to curly leaf (clf-2) mutant and Ler wild-type (WT) plants.**

**Fig. S5 Meristem size and organ initiation analyses of the H3.3 and H3.3K27A lines.**

**Fig. S6 H3.3$^{K27A}$ lines proliferate calli faster, and fail to regenerate shoots on Shoot inducing medium**.

**Fig. S7 Cell elongation and stomata patterning are affected in the H3.3$^{K27A}$ lines.**

**Fig. S8 Additional stem morphology analyses of the H3.3$^{K27A}$ lines: epidermis and cortex.**

**Fig. S9 Additional stem morphology analyses of the H3.3$^{K27A}$ lines: epidermis and cortex layer measurements.**

**Fig. S10 Expression of *WOX4* in the lower inflorescence stems of the H3.3$^{K27A}$ plants.**

**Fig. S11 DNA ploidy levels in H3.3 and H3.3$^{K27A}$ lines.**

**Fig. S12 The H3.3$^{K27A}$ mutant does not induce dramatic changes in chromatin structure (qualitative analyses).**

**Fig. S13 Analysis of H3 mark abundance in H3.3$^{K27A}$ vs H3.3 expressing lines.**

**Fig. S14 GO of genes up-regulated in the H3.3$^{K27A}$ plants and enrichment in transcription factor (TF) classes.**

**Fig. S15 GO of genes down-regulated in the H3.3$^{K27A}$ plants and proportions of transcription factors (TFs) in the various populations of down-regulated genes in overlap with H3K27me3 and or *emf2* targets.**

**Fig. S16 Metabolite profiles of H3.3$^{K27A}$ #B diverge from that of WT and H3.3 #C.**

**Table S1 List and sequences of oligonucleotides used in the study.**

**Table S2 RNA-seq data with the full list of *Arabidopsis thaliana* genes and log2FC between H3.3$^{K27A}$ (#B) and H3.3 (#C) calli.**

**Table S3 Overlap between the lists of H3.3$^{K27A}$ up-regulated genes and H3K27me3 targets or/and *emf2* up-regulated genes.**

**Table S4 Overlap between the lists of H3.3$^{K27A}$ down-regulated genes and H3K27me3 targets or/and *emf2* down-regulated genes.**

**Table S5 Metabolomics data with the full list of *Arabidopsis thaliana* genes and log2FC between H3.3$^{K27A}$ (#B) and H3.3 (#C) lines**.

**Table S6 Metabolomics data for H3.3$^{K27A}$ vs H3.3, mapped to KEGG for the phenylpropanoid biosynthesis pathways**.

**Table S7 Genes mis-regulated in H3.3K27A and downregulated in *wox4* stems.**

**Methods S1 Supporting information for the Material and methods section.**

**Fig. S1 Sequences of the H3.3- and H3.3$^{K27A}$ -encoding transgenes (designed from the *HTR5* sequence) and of the H3.3-encoding *HTR5* (At4g40040).** Exonic regions are in uppercase letters, while intronic regions are in grey lowercase letters. The codon for amino acid 27 of H3.3 is highlighted in yellow. Sequence changes for Lysine to Alanine substitution appear in red. Polymorphic sequences introduced in the 3' end of the transgenes, for differential PCR amplification between transgene and endogene appear in blue. The oMLMc2-oMLMc8 primer pair (sequences underlined) allows amplification of a 159 bp fragment from the transgene, while the oMLMc2-oMLMc11 primer pair (sequences underlined) allows amplification of a 159 bp fragment specific to the endogene. The primer sequences are also given in Table S1.

**Fig. S2 Selection of sets of H3.3 and H3.3$^{K27A}$ lines with comparable transgene expression and analysis of H3 protein accumulation in the soluble vs insoluble fraction. (a-b)** Overview (side and top views) of (a) 25 day-old and (b) 45-day-old plants corresponding to the different lines analysed in (c) for expression of H3.3-coding transgene and genes. Wild-type (WT) non-transgenic plants, transgenic lines expressing a wild-type version of H3.3 (H3.3, lines #C and #A) or a H3.3 variant devoid of Lysine in position 27 (H3.3$^{K27A}$, lines (H3.3, lines #37, #B and #D), all in the Landsberg erecta (Ler) ecotype. **(c)** RT-PCR for expression of

*HTR5* transgenes vs endogenous *HTR5, HTR4* or *HTR8* genes, in *Arabidopsis thaliana* transgenic lines expressing H3.3 and H3.3$^{K27A}$ variants. Polymorphic Sequences (APS), translationally silent, allow to discriminate between endogenous H3 and transgenic H3-derived transcripts, thanks to the use of specific PCR primers. This allowed to perform RT-PCR discriminative for endogenous H3 and transgenic H3-derived transcript detection. 21 cycles of PCR. **(d)** Western-blot for of H3 protein accumulation in soluble and insoluble protein extracts. Soluble and insoluble protein fractions were isolated from 4-d-old Ler wild-type (WT), H3.3 control (H3.3 #A and #C), H3.3$^{K27A}$ (H3.3$^{K27A}$ #B and #37) and *curly leaf* (*clf-2*) plants grown on ½ Murashige and Skoog (MS) medium in a growth chamber under a 16-h light/8-h dark cycle at 22°C. Fractions were analysed by SDS-PAGE and western bloting using the anti-UGPase antibody specific of the soluble fraction and the anti-histone H3 and anti-H3K27me3 antibodies specific of the insoluble/chromatin fraction. Unmethylated H3 proteins were only detected in H3.3 control and H3.3$^{K27A}$ overexpressing lines.

**Fig. S3 H3.3$^{K27A}$ lines flower early in mid-day (MD) and long-day (LD) conditions. Plots, illustrating the number of leaves (a)** and the number of days to bolting **(b)** for the plants of WT, H3.3, H3.3$^{K27A}$ (#B, #37) and and *clf-2,* grown in Short Day (SD, 8h /16h), Mid-Day (MD, 12h/12h) and Continuous Light (CL, 24h light) conditions. The black line in the plots represents the median; the dots are the values of individual samples. Sample size: at least 20 plants for all genotypes, the significance of the observed effect was estimated by the Tukey pairwise comparison test (p-val. < 0.05).

**Fig. S4 Expression analysis of key flowering genes in the H3.3 control (H3.3 #C) and H3.3K27A (H3.3K27A #B and #37) compare to curly leaf (clf-2) mutant and Ler wild-type (WT) plants. (a)** Boxplots showing the relative expression levels of MADS AFFECTING FLOWERING 1/FLOWERING LOCUS M (FLM)/(MAF1), MAF2, MAF3, FLOWERING LOCUS C (FLC) and SUPPRESSOR OF OVEREXPRESSION OF CO1 (SOC1). Expression was measured by quantitative PCR after reverse transcription (RT-qPCR), on 4-day-old seedlings grown on ½ Murashige and Skoog (MS) medium in a growth chamber under a 16-h light/8-h dark cycle at 22°C. **(b)** Boxplots showing the relative expression levels of SOC1 in 10-day-old seedlings. Black lines inside the boxes represent medians. Two independent biological experiments are presented relative to Col-0 levels (set to 1). ACTIN2 and GAPDH were used as reference genes for normalization. Letters indicate significant differences (Student's t-test with Benjamini–Hochberg FDR correction, p-val. < 0.05).

**Fig. S5 Meristem size and organ initiation analyses of the H3.3 and H3.3K27A lines. (a)** Images of the representative samples of dissected shoot apical meristems from the WT, H3.3, H3.3K27A #B, and H3.3K27A #37 plants, taken with KEYENCE VHX microscope, Scale bars images – 50µm. **(b)** Plot, illustrating the radius of shoot apical meristems of the WT, H3.3, H3.3K27A #B, and H3.3K27A #37 plants (as shown in (a)), dissected after the opening of the first flower. Sample size: WT = 24, H3.3 = 27, H3.3K27A #B = 25, and H3.3K27A #37=25, from 2 independently grown populations. **(c)** Representative images of the top view on the inflorescences of WT, H3.3, H3.3K27A #B, and H3.3K27A #37 plants. Scale bars images – 2mm. **(d)** Plot, illustrating the number of flowers, produced on the main stems of plants from the same genotypes; Sample size: WT = 38, H3.3 = 39, H3.3K27A #B = 42, and H3.3K27A #37=44. **(e)** Plot, depicting the number of side branches formed by the WT, H3.3, H3.3K27A #B, and H3.3K27A #37 plants. All measurements were performed on the material from 2 independently grown plant populations (LD, 21°C), the black line in the plots represents the median; the dots are the values of individual samples. Significance of the observed effect was estimated by the Tukey pairwise comparison test (p-val. < 0.05).

**Fig. S6 H3.3$^{K27A}$ lines proliferate calli faster, and fail to regenerate shoots on Shoot inducing medium. (a)** Leaves 3 and 4 of 16 day-old seedlings of designated mutants were cultured on callus inducing media (6 leaves in each replicate). Images were taken on day 30. **(b)** Thirty day-old calli derived from leaves of designated mutants were re-cultured on shoot inducing media (SIM) under light. Images were taken on 9 and 28 days of culturing on SIM. Callus from H3.3$^{K27A}$ lines failed to regenerate shoots. On the middle, higher magnification of the boxed region. On the right: plates of designated lines showing that all calli exhibit similar phenotype. Scale bar = 1 mm.

**Fig. S7 Cell elongation and stomata patterning are affected in the H3.3$^{K27A}$ lines.** Scanning Electron Microscopy (SEM) analysis of stems from 32 day-old plants. WT L*er* ecotype, two independent lines of H3.3 control and H3.3$^{K27A}$ plants. The H3.3$^{K27A}$ lines exhibit aberrant cell shape and stomata distribution. Scale bar =10µm.

**Fig. S8 Additional stem morphology analyses of the H3.3$^{K27A}$ lines: epidermis and cortex.** Representative Images of the transversal microtome sections (8 µm) of paraphrastic-embedded tissues from the base of the main inflorescence stem (0.7 - 1 cm above the rosette and under the first side branch) of WT, H3.3, H3.3$^{K27A}$ #B, and H3.3$^{K27A}$ #37 (2.5 weeks after bolting) plants, stained with Toluidine blue. Scale bar = 100µm.

**Fig. S9 Additional stem morphology analyses of the H3.3$^{K27A}$ lines: epidermis and cortex layer measurements. (a)** Schematic drawings of the transversal sections of Arabidopsis inflorescence stem, where the colored lines indicate the regions taken for the measurements shown in (b), with A and B indicating the shape of a vasculature bundle, C – thickness of the phloem tissue layer and D – thickness of the epidermis and cortex tissue layers. **(b)** Plots depicting the measurements of the shape of the vasculature bundles as a ration between the A and B measurements (as depicted in (a)). Sample sizes: WT = 20, H3.3 = 19, H3.3$^{K27A}$ #B = 28, and H3.3$^{K27A}$ #37=18. The middle panel illustrates the thickness if phloem tissue layer in the stems of WT = 20, H3.3 = 19, H3.3$^{K27A}$ #B = 28, and H3.3$^{K27A}$ #37=18 (n=21, 20, 28 and 19 respectively). The right panel shows the plot of the epidermis and cortex tissue layers thickness in the samples from the same genotypes (n=30, 29, 26 and 41 respectively). The measurements for each phenotype were taken on at least 4 plants from 2 independently grown populations. The black line in the plots represents the median; the dots are the values of individual measurements, the letters indicate the statistical groups, established with the by the Tukey pairwise comparison test (p-val. < 0.05).

**Fig. S10 Expression of *WOX4* in the lower inflorescence stems of the H3.3$^{K27A}$ plants.** Histograms illustrating the abundance of the *WOX4* transcript in the base parts of inflorescence stems of the H3.3, WT, H3.3$^{K27A}$ #B and H3.3$^{K27A}$ #37 plants, measured by RT-qPCR. The data is normalised to *TUB4* and represented as a fold change over the control. The error bars indicate the standard deviation of the 3 technical repeats, the data from two independent biological replicates is shown side by side **(a,b)**. Letters indicate significant differences (Student's *t*-test with Benjamini–Hochberg FDR correction, p-val. < 0.05).

**Fig. S11 DNA ploidy levels in H3.3 and H3.3$^{K27A}$ lines.** Ploidy levels were determined by flow cytometry on nuclei extracted from roots of 7 day-old seedlings grown on ½ MS medium in a growth chamber under LD (16h light/8h dark) at 22°C. Data shown are mean±SEM (n=15,000), based on technical duplicate.

**Fig. S12 The H3.3$^{K27A}$ mutant does not induce dramatic changes in chromatin structure (qualitative analyses).** Immuno-localization of different histone marks on Arabidopsis root tip. Paraformaldehyde-fixed nuclei from 7 day-old *in vitro*-grown seedlings were used. Left panel shows representative Z-stack projection images of H3K9me2 (green) and H3K27me1 (red) distributions in the nuclei with the DAPI staining (blue). Right panel shows representative DAPI (blue), H3K4me3 (green) and

H3K27me3 (red) distributions in the nuclei with the DAPI staining (blue). The DNA At least 100 nuclei from different experiments were observed for each Arabidopsis line. Scale bars for all images: 2µm.

**Fig. S13 Analysis of H3 mark abundance in H3.3$^{K27A}$ vs H3.3 expressing lines.** Western blot analysis of histone lysine methylation and acetylation in 4-d-old Ler wild-type (WT), H3.3 control (H3.3 #A and #C), H3.3$^{K27A}$ (H3.3$^{K27A}$ #B and #37) and curly leaf (clf-2) plants. Nuclear protein extracts from 4-d-old WT, overexpressing and T-DNA mutant plants, grown on ½ Murashige and Skoog (MS) medium in a growth chamber under a 16-h light/8-h dark cycle at 22°C, were analyzed using antibodies specific for modified and non-modified histone H3.

**Fig. S14 GO of genes up-regulated in the H3.3$^{K27A}$ plants and enrichment in transcription factor (TF) classes (log2FC$\geq$1, p-value < 0.05),** for **(a)** the 534 genes that also are targets of H3K27me3; for **(b)** the 109 genes also up-regulated in and *emf2* (log2FC$\geq$1) and targets of H3K27me3.

**Fig. S15 GO of genes down-regulated in the H3.3$^{K27A}$ plants and proportions of transcription factors (TFs) in the various populations of down-regulated genes in overlap with H3K27me3 and or *emf2* targets.** GO of genes down-regulated in the H3.3$^{K27A}$ plants (log2FC≤-1, p-value < 0,05) for **(a)** the 86 genes that also are targets of H3K27me3; for **(b)** the 18 genes that are also down-regulated in *emf2* (log2FC≤-1) and targets of H3K27me3. **(c)** Percentages of TFs in the various populations of down-regulated genes in overlap with H3K27me3 targets and/or *emf2* up-regulated genes. The full list of corresponding genes can be found in the Suppl. Table S4.

**Fig. S16 Metabolite profiles of H3.3$^{K27A}$ #B diverge from that of WT and H3.3 #C. (a)** Loading plot of metabolite profiling data of WT, H3.3 #C and H3.3$^{K27A}$ #B. Each point represents a metabolite. The loading plot was constructed using MetaboAnalyst 5.0. Data were analysed by range scaling. **(b-e)** Volcano plots showing the differential metabolites between (b) H3.3$^{K27A}$ #B and H3.3 #C or (c) H3.3$^{K27A}$ #37 and H3.3$^{K27A}$ #A, (d) H3.3 #A and WT, or (e) H3.3 #C and WT. Each point in the volcano plot represents one metabolite (or bucket). Significant buckets were calculated with a fold change (FC) threshold of +/- 2 and a minimum p value of 0.05 (indicated in orange).

**Table S1 List and sequences of oligonucleotides used in the study.**

**Table S2 RNA-seq data with the full list of *Arabidopsis thaliana* genes and log2FC between $H3.3^{K27A}$ (#B) and H3.3 (#C) calli**. Columns D-K show mRNA levels (in RPKM: Reads Per Kilobase of transcripts per Million mapped reads) of callus derived from cotyledons, in three biological replicates, for the $H3.3^{K27A}$ #B and H3.3#C lines. Columns G and K give the mean from the 3 replicates. Column L gives the log2 Fold Change between for $H3.3^{K27A}$ #B compared to H3.3#C. All *Arabidopsis thaliana* genes are shown, with rows sorted by gene ID number.

**Table S3 Overlap between the lists of $H3.3^{K27A}$ up-regulated genes and H3K27me3 targets or/and *emf2* up-regulated genes.** Ten sheets are displayed, corresponding to the different gene populations represented in Fig. 3b, for up-regulated genes (log2FC≥1). In the first sheet, corresponding to all 1764 genes up-regulated in $H3.3^{K27A}$ #B as compared to H3.3#C, Columns E-L show mRNA levels (in RPKM: Reads Per Kilobase of transcripts per Million mapped reads) of callus derived from cotyledons, in three biological replicates, for the $H3.3^{K27A}$ #B and H3.3#C lines. Columns H and L give the mean from the 3 replicates. Column M gives the log2 Fold Change between for $H3.3^{K27A}$ #B compared to H3.3#C. Rows are sorted by TF family, and then gene ID number. The last sheet displays the Representation factors and p-values for statistical significance of the overlaps between the groups of genes (H3.3K27A up-regulated AND H3K27me3 targets; H3.3K27A up-regulated AND *emf2* up-regulated).

**Table S4 Overlap between the lists of $H3.3^{K27A}$ down-regulated genes and H3K27me3 targets or/and *emf2* down-regulated genes.** Ten sheets are displayed, corresponding to the different gene populations represented in Fig. 3c, for down-regulated genes (log2FC ≤-1). In the first sheet, corresponding to all 1337 genes down-regulated in $H3.3^{K27A}$ #B as compared to H3.3#C, Columns E-L show transcript levels (in RPKM: Reads Per Kilobase of transcripts per Million mapped reads) of callus derived from cotyledons, in three biological replicates, for the $H3.3^{K27A}$ #B and H3.3#C lines. Columns H and L give the mean from the 3 replicates. Column M gives the log2 Fold Change between for $H3.3^{K27A}$ #B compared to H3.3#C. Rows are sorted by TF family, and then gene ID number. The last sheet displays the Representation factors and p-values for statistical significance of the overlaps between the groups of genes (H3.3K27A down-regulated AND H3K27me3 targets; H3.3K27A down-regulated AND *emf2* down-regulated).

**Table S5 Metabolomics data with the full list of *Arabidopsis thaliana* genes and fold changes (FC) between H3.3$^{K27A}$ (#B) and H3.3 (#C) lines**. Eight sheets are displayed, corresponding to the Differential data grouped as color groups from Kegg Mapper, the color keys for Kegg Mapper, and the analyte lists as given by Phenol Explorer (http://phenol-explorer.eu/), LipidMaps (https://www.lipidmaps.org/), PlantCyc (https://plantcyc.org/), Knapsack (http://www.knapsackfamily.com/); the last sheet corresponds to the Raw data.

**Table S6 Metabolomics data for H3.3$^{K27A}$ vs H3.3, mapped to KEGG for the phenylpropanoid biosynthesis pathways**. Three sheets are displayed, corresponding to the initial data set as annotated from PhenolExplorer, and to the analytes that are over-represented (22) and those that are under-represented (60) in H3.3$^{K27A}$ #B as compared to H3.3#C (p-val. < 0.05). Rows are sorted by the KEGG pathway.

**Table S7 Genes mis-regulated in H3.3K27A and downregulated in *wox4* stems.** A single sheet displays the lists of genes that are upregulated in H3.3$^{K27A}$ #B (column A), downregulated in H3.3$^{K27A}$ #B (column B), and downregulated in stems of *wox4* plants (column C). The data on the right represents the common genes among those upregulated in H3.3$^{K27A}$ #B and downregulated in *wox4* (columns H to O), and those that are downregulated in H3.3$^{K27A}$ #B and downregulated in *wox4* (columns R to Y). In each case, the highlighted columns (N-O and X-Y) represent the genes which expression is enriched in xylem (columns N and X) and those that are associated with H3K27me3-enriched regions. Columns K and U give the log2 Fold change for expression. This table reveals a strong enrichment in genes mis-regulated in H3.3$^{K27A}$ among those that are WOX4 targets (Representation factor: 1.8, p-val. < 6.696e-06).

**Methods S1 Supporting information for the Material and methods section.**

*Construction and selection of the H3.3 variant lines*

Plant transformation and primary transformant selection

*Arabidopsis thaliana* Ler plants were used. Primary transformants, resistant to kanamycin, were verified by PCR using the primer pairs described in Fig. S1 and Table S1. Lines with a single insertion locus were brought to the T3 generation for transgene expression analyses.

Transgene expression analyses by RT-PCR

For transgene expression by RT-PCR, total RNA was isolated from 8-days-old seedling growing on MS plate using the Qiagen RNeasy Plant Mini Kit (Cat. No. / ID: 74904). After a DNase treatment (ezDNase SuperScript IV VILO, ThermoFisher, Cat. No. 11756050), first strand cDNA synthesis was performed with 2µg of total RNA using SuperScript IV VILO, ThermoFisher, Cat. No. 11756050. For each PCR reaction, 1/40$^{th}$ of the synthesized cDNA were used. The annealing temperature was 55°C for all primer pairs (described in Table S1) and 25 cycles of PCR were performed for *HTR5* transgene, *HTR4* and *HTR8* endogenes, 21 cycles for *EF1alpha* and *HTR5* endogene.

in cyto *analyses and Immuno-detection*

Seedlings were fixed for 30 minutes at 4°C in 4% (w/v) paraformaldehyde in MTSB buffer (5 mM EGTA, 2 mM MgCl2, 50 mM PIPES, pH 6.9). Root tips were digested for 10 min at 37°C in a enzymatic mixture (2.5% cellulase, 2.5% pectinase, 2.5% pectolyase (w/v) in MTSB) and squashed. Immunostaining was performed as described (Batzenschlager *et al.*, 2015). The primary antibodies anti-H3K27me3 (Diagenode, lot n° A932-00234P, 1/500), anti-H3K27me1 (Diagenode, lot n°A1818P, 1/1000), anti-H3K4me3 (Diagenode, lot n°001-13, 1/500) and anti-H3K9me2 (Diagenode, lot n°-001-12, 1/500) were incubated at 4°C. Signals were revealed with the following secondary antibodies: Alexa 488–conjugated anti-mouse IgG (1/200) or Alexa 568–conjugated goat anti-rabbit IgG (1/300) from Molecular Probes® (Life Technologies). Root tips were mounted in Vectashield (Vector Laboratories) containing 2 µg/ml DAPI.

*Ploidy Level Measurement*

Nuclear DNA content was measured using the CyStain UV Precise P Kit (Partec) according to the manufacturer's instructions. Nuclei from the root system of 7 DAS seedlings grown on ½ MS medium in a growth chamber under a 16-h light/8-h dark cycle at 22°C, were released in nuclei extraction buffer (Partec) by lightly chopping the material with a razor blade, stained with 4',6-diamidino-2-phenylindole buffer, and filtered through a Celltrics 30-µm mesh (Partec). Approximatively 15000 isolated nuclei were used for each analysed line, using the Attune Cytometer and the Attune Cytometer software (Life Technologies) recording the relative fluorescence intensities. Flow cytometry experiments were repeated two times.

*Protein extraction and immunoblotting*

Fractions of soluble/insoluble proteins were prepared as previously described (Schalk *et al.*, 2017). Briefly, after liquid nitrogen grinding, aerial parts from 4-day-old seedlings were incubated at 4°C during

30 min on a rotating wheel in a lysis buffer (25 mM Tris·HCl pH 8.0, 0.3 M NaCl, 1 mM EDTA, 10% (vol/vol) glycerol, 1% (vol/vol) Nonidet P-40, 0.2 mM phenylmethylsulfonyl fluoride, and EDTA-free protease inhibitor cocktail (1 tablet/50 mL; Roche)). Cellular debris were then removed by Miracloth (Calbiochem) filtering and centrifugation (2,000 x g, 5 min, 4°C). The insoluble fraction was separated from the soluble one by centrifugation (13,000 x g, 10 min, 4°C), resuspended in the lysis buffer and used for immunoblotting.

Nuclear protein extracts for histone modification analyses were prepared as described previously (Zhang *et al.*, 2020). Briefly, aerial parts from 4-day-old seedlings were grinded into a fine powder under liquid nitrogen and incubated at 4°C during 20 min on a rotating wheel in a lysis buffer (50 mM HEPES pH7.5, 0.5 M NaCl, 50 mMEDTA, 1% (vol/vol) TritonX-100, 10% (vol/vol) glycerol, 0,04% (vol/vol) β-mercaptoethanol, and EDTA-free protease inhibitor cocktail (1 tablet/50 mL; Roche)). After filtration through Miracloth (Calbiochem) and centrifugation (2,000 x g, 20 min at 4°C), the pellet was resuspended in 1 x SDS-PAGE sample buffer and used for immunoblotting.

*RNA-seq raw-reads treatment and analysis*

Raw-reads were subjected to a filtering and cleaning procedure. The Trimmomatic tool (Bolger *et al.*, 2014) was used to remove Illumina adapters from the reads. Next, the FASTX Toolkit (http://hannonlab.cshl.edu/fastx_toolkit/index.html, version 0.0.13.2) was used to trim read-end nucleotides with quality scores <30, using the FASTQ Quality Trimmer, and to remove reads with less than 70% base pairs with a quality score ≤30 using the FASTQ Quality Filter. Cleaned reads were mapped to the reference genome of *Arabidopsis thaliana* (TAIR10) using TopHat2 (Kim *et al.*, 2013) with an average mapping rate of 90.50%. Gene abundance estimation was performed using Cufflinks v. 2.2 (Trapnell *et al.*, 2010). PCA analysis and Heatmap visualization were performed using R Bioconductor (Gentleman *et al.*, 2004). Gene expression values were computed as FPKM. Differential expression analysis was completed using the DESeq2 R package (Love *et al.*, 2014).

*Mass-spectrometry for non-targeted metabolomic analyses*

For metabolite identification, 300 mg of fresh leaves powder from each genotype were resuspended in 1.5ml cold (5°C) methanol spiked with an internal standard of deuterium labelled $[^2H_6]$(+)-cis,trans-abscisic acid ($^2H_6$-ABA) at 0.1µg/ml. After 10 seconds of vortexing, the samples were stored for 16h at -20°C, then centrifuged at 13000 rpm for 10 min at 4°C. The supernatant was collected and dried by sublimation using a SpeedVac concentrator (Savant SPD121P, Thermo Fisher). The pellet was resuspended in 1.5mL cold MeOH, shaken for 10 minutes, then centrifuged at 13000 rpm for 10 min at 4°C, and the supernatant was collected to be dried in the same glass vial as the first extraction. A third extraction was performed as the second extraction. The dried supernatant from the 3 successive extractions was resuspended in 300µL of MeOH for analysis in liquid chromatography coupled to high resolution mass spectrometry (LC-HRMS).

Then, samples were analyzed using liquid chromatography coupled to high resolution mass spectrometry on an UltiMate 3000 system (Thermo) coupled to an Impact II (Bruker) quadrupole time-of-flight (Q-TOF) spectrometer. Chromatographic separation was performed on an Acquity UPLC ® BEH C18 column (2.1x100mm, 1.7µm, Waters) equipped with and Acquity UPLC ® BEH C18 pre-column (2.1x5mm, 1.7µm, Waters) using a gradient of solvents A (Water, 0.1% formic acid) and B (MeOH, 0.1% formic acid). Chromatography was carried out at 35°C with a flux of 0.3mL.min$^{-1}$, starting with 5% B for 2 minutes, reaching 100% B at 10 minutes, holding 100% for 3 minutes and coming back to the initial condition of 5% B in 2 minutes, for a total run time of 15 minutes. Samples were kept at 4°C, 3µL were injected in full loop mode with a washing step after sample injection with 150µL of wash solution ($H_2O$/MeOH, 90/10, v/v). The spectrometer was equipped with an electrospray ionization (ESI) source and operated in positive ion mode on a mass range from 20 to 1000 Da with a spectra rate of 2Hz in AutoMS/MS fragmentation mode. The end plate offset was set at 500 V, capillary voltage at 2500 V, nebulizer at 2 Bar, dry gas at 8 L.min$^{-1}$ and dry temperature at 200°C. The transfer time was set at 20-70µs and MS/MS collision energy at 80-120% with a timing of 50-50% for both parameters. The MS/MS cycle time was set to 3 seconds, absolute threshold to 816 cts and active exclusion was used with an exclusion threshold at 3 spectra, release after 1 min and precursor ion was reconsidered if the ratio current intensity/previous intensity was higher than 5. A calibration segment was included at the beginning of the runs allowing the injection of a calibration solution from 0.05 to 0.25min. The calibration solution used was a fresh mix of 50mL isopropanol/water (50/50, v/v), 500µL NaOH 1M, 75µL acetic acid and 25µL formic acid. The spectrometer was calibrated on the $[M+H]^+$ form of reference ions (positif : 57 masses from m/z 22.9892 to m/z 990.9196) in high precision calibration (HPC) mode with a standard deviation below 1ppm before the injections for each polarity mode, and re-calibration of each raw data was performed after injection using the calibration segment. Raw data were processed in MetaboScape 4.0 software (Bruker): molecular features were considered and grouped into buckets containing one or several adducts and isotopes from the detected ions with their retention time and MS/MS information when available. The parameters used for bucketing are a minimum intensity threshold of 10000, a minimum peak length of 4 spectra, a signal-to-noise ratio (S/N) of 3 and a correlation coefficient threshold set at 0.8. The $[M+H]^+$ ion was authorized as primary ion,

$[M+Na]^+$, $[M+NH4]^+$, $[M+K]^+$ and $[M-H_2O+H]^+$ were authorized as seed ions. Replicate samples were grouped and only the buckets found in 80% of the samples of one group were extracted from the raw data. The obtained list of buckets was annotated using SmartFormula to generate raw formula based on the exact mass of the primary ions and the isotopic pattern. The maximum allowed variation on the mass (□m/z) was set to 3ppm, and the maximum mSigma value (assessing the good fitting of isotopic patterns) was set to 30. To put a name on the obtained formulae, analyte lists were derived from KNApSAcK (http://www.knapsackfamily.com/), PlantCyc (https://plantcyc.org/), PhenolExplorer (http://phenol-explorer.eu/), LipidMaps (https://www.lipidmaps.org/) and SwissLipids (https://www.swisslipids.org/). The parameters used for the annotation with the analyte lists are the same as for SmartFormula annotation. Stastitical analysis was performed in MetaboScape 4.0 with 8 samples per group using the areas of the peaks as the unit of reference. To comply with the small number of samples, a Wilcoxon rank-sum test was used to compare the groups to each other. Compounds were considered as statistically differential between two groups using the thresholds of $p$-value ≤ 0.05 and fold change ≥ 2 or ≤ -2. The $p$-values and fold changes from the tests were used for the chemical enrichment analysis in ChemRICH, keeping the differential and non-differential compounds to allow the enrichment analysis. A Principal component analysis (PCA) was applied to assess metabolomic data and to provide an overview of variation in datasets, using MetaboAnalyst 5.0. Volcano plots based on a non-parametric Mann-Whitney t-test statistics were used to measure differentially accumulated metabolites and fold changes simultaneously.

Batzenschlager M, Lermontova I, Schubert V, Fuchs J, Berr A, Koini MA, Houlné G, Herzog E, Rutten T, Alioua A, *et al.* 2015. Arabidopsis MZT1 homologs GIP1 and GIP2 are essential for centromere architecture. *Proceedings of the National Academy of Sciences of the United States of America* 112: 8656–8660.

Bolger AM, Lohse M, Usadel B. 2014. Trimmomatic: a flexible trimmer for Illumina sequence data. *Bioinformatics* 30: 2114–2120.

Gentleman RC, Carey VJ, Bates DM, Bolstad B, Dettling M, Dudoit S, Ellis B, Gautier L, Ge Y, Gentry J, *et al.* 2004. Bioconductor: open software development for computational biology and bioinformatics. *Genome Biology*: 16.

Kim D, Pertea G, Trapnell C, Pimentel H, Kelley R, Salzberg SL. 2013. TopHat2: accurate alignment of transcriptomes in the presence of insertions, deletions and gene fusions. *Genome Biology* 14: R36.

Love MI, Huber W, Anders S. 2014. Moderated estimation of fold change and dispersion for RNA-seq data with DESeq2. *Genome Biology* 15: 550.

Schalk C, Cognat V, Graindorge S, Vincent T, Voinnet O, Molinier J. 2017. Small RNA-mediated repair of UV-induced DNA lesions by the DNA DAMAGE-BINDING PROTEIN 2 and ARGONAUTE 1. *Proceedings of the National Academy of Sciences* 114.

Trapnell C, Williams BA, Pertea G, Mortazavi A, Kwan G, van Baren MJ, Salzberg SL, Wold BJ, Pachter L. 2010. Transcript assembly and quantification by RNA-Seq reveals unannotated transcripts and isoform switching during cell differentiation. *Nature Biotechnology* 28: 511–515.

Zhang X, Ménard R, Li Y, Coruzzi GM, Heitz T, Shen W-H, Berr A. 2020. Arabidopsis SDG8 Potentiates the Sustainable Transcriptional Induction of the Pathogenesis-Related Genes PR1 and PR2 During Plant Defense Response. *Frontiers in Plant Science* 11: 277.
